# Deletion of a non-canonical regulatory sequence causes loss of *Scn1a* expression and epileptic phenotypes in mice

**DOI:** 10.1101/766634

**Authors:** Jessica L. Haigh, Anna Adhikari, Nycole A. Copping, Tyler Stradleigh, A. Ayanna Wade, Rinaldo Catta-Preta, Linda Su-Feher, Iva Zdilar, Sarah Morse, Timothy A Fenton, Anh Nguyen, Diana Quintero, Samrawit Agezew, Michael Sramek, Ellie J Kreun, Jasmine Carter, Andrea Gompers, Jason Lambert, Cesar P. Canales, Len A. Pennacchio, Axel Visel, Diane E. Dickel, Jill L. Silverman, Alex S. Nord

**Author notes:** These authors contributed equally (Jessica L. Haigh, Anna Adhikari). These authors are co-corresponding (Jill L. Silverman, Alex S. Nord). Corresponding author emails: Alex S. Nord and Jill Silverman.

## Abstract

Genes with multiple co-active promoters appear common in brain, yet little is known about functional requirements for these potentially redundant genomic regulatory elements. *SCN1A,* which encodes the Na_V_1.1 sodium channel alpha subunit, is one such gene with two co-active promoters. Mutations in *SCN1A* are associated with epilepsy, including Dravet Syndrome (DS). The majority of DS patients harbor coding mutations causing *SCN1A* haploinsufficiency, however putative causal non-coding promoter mutations have been identified. To determine the functional role of one of these potentially redundant *Scn1a* promoters, we focused on the non-coding *Scn1a* 1b regulatory region, previously described as a non-canonical alternative transcriptional start site. Mice harboring a deletion of the extended evolutionarily-conserved 1b non-coding interval exhibited surprisingly severe reductions of *Scn1a* and Na_V_1.1 expression in brain with accompanying electroencephalographic seizures and behavioral deficits. This work identified the 1b region as a critical disease-relevant regulatory element and provides evidence that non-canonical and seemingly redundant promoters can have essential function.

## Introduction

A large proportion of brain-expressed and, indeed, all mammalian genes are believed to rely on multiple alternative promoters^1–3^. For many genes, the alternative promoters produce distinct 5’ untranslated regions but otherwise similar mRNA products, leading to identical proteins from distinct transcription start sites (TSSs)^4,5^. Much of the focus on understanding the role of alternative promoters in mammalian transcriptional regulation has been on the potential for discrete function via producing specific isoforms or compartmentalized expression in specific cells or tissues^6–9^. However, TSS activity mapping has found many genes where alternative promoters are also active in the same tissue^10,11^. More recent evidence from single cell RNA sequencing and chromosome conformation suggests that annotated alternative promoters are frequently co-active in the same cells and physically interact in the nucleus^12–14^. However, in contrast to work on the requirement for alternative promoters with presumed discrete activity, studies investigating the functional requirement for apparently redundant co-active promoters are lacking.

Epilepsy is one of the most common neurological disorders, with both rare highly-penetrant and common variants contributing to genetic etiology. Mutations in *SCN1A,* which encodes the Na_V_1.1 sodium channel alpha subunit, result in a range of epilepsy phenotypes from generalized febrile seizures to Dravet Syndrome (DS), a severe childhood-onset disorder^15–17^. The majority of DS cases are caused by heterozygous *de novo* mutations in *SCN1A* resulting in truncation of the protein, with haploinsufficiency of Na_V_1.1 presumed to underlie pathology^18,19^. Mouse models with heterozygous coding mutations in *Scn1a* recapitulate features of DS, including seizures and sudden unexpected death in epilepsy (SUDEP)^20–26^. DS remains pharmacoresistant, with generalized tonic-clonic seizures beginning in the first year of life and common comorbid neurodevelopmental disorder (NDD) behavioral phenotypes including cognitive impairments and ataxia^27–29^.

*SCN1A* transcripts have a variable 5’ untranslated region (UTR) containing one of two TSSs, 1a and 1b, that are conserved between human and mouse^30,31^. The proteins produced from 1a and 1b are expected to be identical. 1a was found to be the majority TSS for *SCN1A* transcripts across human and mouse (54% and 52% RACE transcripts, respectively). No strong region-specific differences in 1a versus 1b TSS usage across brain regions have been identified in previous work^30,32^. 1a (also referred to as h1u) has been defined as the major *SCN1A* promoter, however, comparison across brain tissues in human and mouse suggests that 1a and 1b are co-active, with ∼35% of transcripts arising from 1b^30^. The apparent functional redundancy of 1a and 1b promoter activity and 1a- and 1b-associated *SCN1A* transcripts raises the question of whether there are distinct roles or requirements of the 1a and 1b UTR and regulatory DNA sequences.

In addition to serving as an example in which to dissect the role of multiple co-active promoters, there is significant disease relevance for understanding the functional requirements for *SCN1A* regulatory DNA elements. *SCN1A* is one of the most common and well-documented genes associated with severe medical consequences of haploinsufficiency. Further, genome-wide association studies (GWAS) have implicated non-coding *SCN1A* DNA variants as contributing to epilepsy risk^33,34^, presumably via more subtle perturbation to transcriptional regulation, and non-coding promoter deletions have been found in DS patients^35,36^. A recent study of common variation in the promoter regions of *SCN1A* found that promoter variant haplotypes reduced luciferase in cells and that such non-coding variants in the functional *SCN1A* allele may modify DS severity^37^. Based on these findings, it is plausible that pathogenic variation in regulatory regions modulates *SCN1A* transcription, contributing to epilepsy. Functional studies are needed to determine the consequences of perturbations to *SCN1A* expression caused by mutations in non-coding DNA.

Here, we investigate *Scn1a* as a model for examining transcriptional and phenotypic consequences associated with loss of a potentially redundant co-active promoter. Combining genomics, neuropathology, behavior, seizure susceptibility, and EEG, we show that the *Scn1a* 1b non-canonical promoter and flanking conserved non-coding DNA sequence is independently essential for expression and brain function via characterizing impact 1b^+/−^ and 1b^−/−^ mice. In addition to mapping an essential regulatory region of a critical disease-relevant gene, our findings provide evidence that non-canonical promoters can play essential roles in general transcriptional activation.

## Results

### Scn1a 1a and 1b chromosomal regions physically interact and share chromatin signatures indicating pan-neuronal transcriptional activator regulatory activity

To define the regulatory landscape of the *SCN1A* locus, we examined publicly-available chromosome conformation (Hi-C), transcriptomic and epigenomic data obtained from analysis of human and mouse brain tissues (**Fig. 1a, d, S1**). We generated contact heatmaps from published Hi-C from prefrontal cortex^38^ (PFC) at 10-kb resolution (**Fig. S1a**), and for additional tissues at 40-kb resolution^38^ (**Fig. S1c**). In PFC, *SCN1A* was located at the boundary of two major TADs (Topological Associated Domains), with extensive local interactions within the *SCN1A* locus. Differential analysis of Hi-C from PFC versus lung showed stronger local interactions in PFC (**Fig. S1b**), while there were no major differences between the PFC and hippocampus, suggesting brain-specific local *SCN1A* chromosomal interactions (**Fig. S1c**).

**Figure 1:**
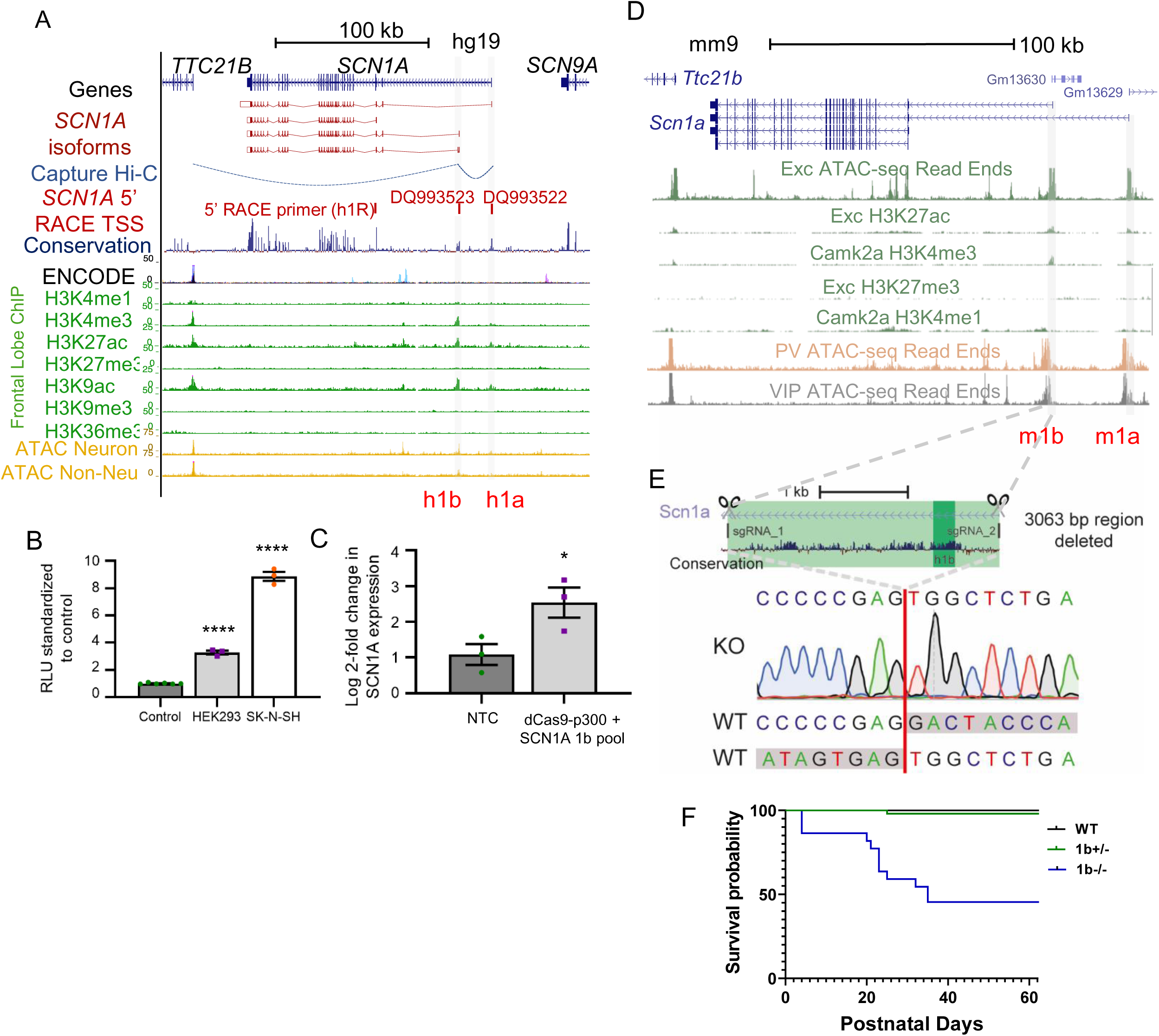
Genomic context of *SCN1A* gene and Scn1a 1b deletion mouse model generation. (a) Human (hg19) *SCN1A* locus showing signal for histone PTMs and ATAC-seq for neuronal and non-neuronal cells derived from dorsolateral PFC. (b) Activity of human 1b region in luciferase assay with minimal promoter in HEK293 (****P < 0.0001) and SK-N-SH cells (****P < 0.0001) shown as mean ± SEM. (c) Transcriptional activation of *SCN1A* using gRNAs targeting 1b co-transfected with dCas9-p300 in HEK293 cells increases *SCN1A* expression (*P = 0.047) when compared to empty vector (EV) as measured by qPCR and normalized to non-transfected control, shown as mean ± SEM. (d) Mouse (mm10) *Scn1a* locus showing signal for histone PTMs and ATAC-seq for neuronal cell types. (e) Mouse Scn1a 1b locus showing guideRNA sequence targets for Cas9-directed deletion of mouse 1b removal of entire 3063 bp conserved region and sequence trace validating deleted region. (f) Survival curve for offspring from 1b^+/−^ by 1b^+/−^ breeding pairs. Data is from 13 litters that dropped from 3 generations of pairings combined. 1b^+/−^ and 1b^+/+^ Logrank p value = 0.5455; 1b^+/+^ versus 1b^−/−^ Logrank p value < 0.0001. For panels a and d see text for data sources.

Previous work using 5’ RACE^30^ and luciferase assays defined regulatory sequences at *SCN1A*, including two genomic intervals, non-coding exons 1a and 1b, that are conserved between human and mouse and where the majority of *SCN1A* transcripts originate^30^ (**Fig. 1a**). The locations of the 1a (GenBank: DQ993522) and 1b (GenBank: DQ993523) TSSs are shown as mapped in earlier work^30,32^. DNA sequence at 1a and 1b is highly evolutionarily-conserved across vertebrates. Notably, conservation at 1b extends nearly 3 kb downstream of the defined UTR transcribed sequence. Interaction models from an independent capture Hi-C dataset^38^ also suggested physical interaction between 1a and 1b as well as between 1b and the nearby *TTC21B* promoter (**Fig. 1a**).

We examined chromatin state at the *SCN1A* locus across seven histone post-translational modifications (PTMs) from human mid frontal lobe^39^ (**Fig. 1a**). The strongest chromatin signatures for regulatory elements were at the previously-defined 1a and 1b loci, with H3K27ac, H3K4me3 and H3K9ac, weak H3K4me1, and absence of H3K27me3, H3k9me3, H3K36me3 in these regions. In ATAC-seq and H3K27ac across the majority of non-CNS tissues profiled in the ENCODE or Roadmap projects, 1a and 1b show reduced or absent signal, further indicating primary importance in the nervous system (data not shown). In addition to 1a and 1b, there were several other non-coding regions showing weaker, but still significant enrichment for H3K27ac in brain, representing potential additional *SCN1A* regulatory elements. ATAC-seq from neuronal and non-neuronal cells from dorsolateral PFC (DLPFC)^40^ showed that neuronal cells have increased chromatin accessibility across *SCN1A* generally (**Fig. 1a**), with specific enrichment at 1a, 1b, and a third region also within the first intron of *SCN1A*. Comparing human neuronal data with mouse ATAC-seq and histone PTM data^41^, accessibility and chromatin states appeared largely conserved (**Fig. 1a** and **1d**). Finally, ATAC-seq data^41^ from sorted neuronal subtypes in mouse, including excitatory neurons and parvalbumin (PV) and vasoactive intestinal peptide (VIP) interneurons, indicated consistent open chromatin at *Scn1a* 1a and 1b across neuron types (**Fig. 1d**).

### The evolutionarily-conserved Scn1a 1b non-coding region acts as a Scn1a transcriptional activator and is essential for survival

Taken together, the comparative and functional genomics data indicates evolutionarily conserved brain-specific pan-neuronal regulatory and TSS activity of 1a and 1b, with evidence for chromosomal physical interaction between the two promoters. The 1b region has been annotated as an alternative TSS, yet the extended region surrounding the annotated transcribed UTR also shows the strongest enrichment for evolutionary conservation and for chromatin signatures associated with strong enhancer activity (i.e. H3K27ac and H3K4me3) across non-coding regions of the *SCN1A* locus. We sought to validate the specific role of 1b DNA in activation of *SCN1A* expression. We used luciferase assay to functionally test the core human 1b (h1b) region in cell lines. A 941-bp region containing 1b and conserved flanking sequence induced expression in HEK293 and SK-N-SH cells when cloned into a vector with a minimal promoter (**Fig. 1b**). To further demonstrate the regulatory role of 1b in *SCN1A* expression, we showed that a pool of 6 sgRNAs targeted to human 1b sequence and delivered along with dCas9-p300, an inactivated Cas9-histone acetyltransferase fusion protein^42^, was sufficient to induce *SCN1A* expression 2.5-fold in HEK293 cells compared to non-transfected control (**Fig. 1c**).

The strength of evolutionary conservation and transcriptional activation-associated epigenomic signatures at the extended 1b interval is paradoxical considering its presumed role as a secondary TSS. Thus, we sought to test whether the extended 1b regulatory region is essential for *Scn1a* expression, and whether loss of this element is sufficient produce seizures and DS-relevant phenotypes in mice. We used CRISPR/Cas9 targeting of C57BL/6N oocytes to generate mice harboring a 3063 bp deletion of the interval flanking the 1b regulatory element of *Scn1a*, removing the entire mammalian conserved region (**Fig. 1e**). We expanded this *Scn1a 1b* deletion line (hereafter referred to as 1b) via at least six generations of breeding to wildtype C57BL/6N (WT) mice, eliminating potential off-target Cas9-induced mutations.

Previous studies have found that mice harboring homozygous coding mutations to *Scn1a* die in the third postnatal week and mice with heterozygous coding mutation exhibit reduced survival^20,26^. In comparison, 46 female WT by heterozygous male (1b^+/−^) harem trio pairings yielded 41 litters and survival rates of 1b^+/−^ and WT littermate pups were indistinguishable (Log-rank p=0.8458), however, female 1b^+/−^ by 1b^+/−^ male harem trios required nearly double the number of pairings (n = 74) to produce only 13 litters. This was caused by both reduced rates of pregnancy, as determined via regularly weighing females, and by increased litter cannibalism in the neonatal period. Among the 13 litters that were produced, the three genotypes were born at expected Mendelian frequencies (**Table S1**). While survival rates for 1b^+/−^ and WT littermate pups from these litters were indistinguishable (Log-rank p=0.5455), 48% of homozygous 1b deletion (1b^−/−^) mice died by weaning (**Fig. 1f**, Logrank p<0.0001). Thus, 1b^+/−^ mice survive, but female carriers were less efficient at producing viable litters, severely impacting generation of homozygous 1b deletion offspring. 1b^−/−^ pups were visibly smaller and failed to thrive, though around half survived to maturity. In addition, 1b^−/−^ exhibited spontaneous seizures during routine handling, consistent with expected neurological impact of significant decrease in *Scn1a* expression.

We tested 1b^+/−^ mice for measures of general health and utilized a Fox developmental battery^43^ and found no deficits in growth, reflexes, and limb strength (**Table S2**). 1b^−/−^ pups were not evaluated for these milestones, as spontaneous seizures were observed, so handling and was minimized to increase survival.

### Loss of extended 1b interval causes loss of Na_V_1.1 across postnatal brain regions

We first sought to test if 1b ablation resulted in changes in amount and regional distribution of *Scn1a* transcript and Na_V_1.1 protein in mouse brain. Expression of *Scn1a*/Na_V_1.1 begins in the early postnatal period and reaches high expression throughout the brain by four weeks of age^44^. We focused on the cerebellum, hippocampus and cortex, where we detected Na_V_1.1 by immunohistochemistry in WT mice by P28 (**Fig. 2a-c**). We tested for reduced *Scn1a* RNA expression via quantitative reverse-transcription PCR (qRT-PCR) performed on cortex, hippocampus and cerebellum of 3-month-old *Scn1a* 1b deletion carriers and WT littermates (WT n=4, 1b^+/−^ n=5, 1b^−/−^ n=7) (**Fig. 2d**). In agreement with GTEx^32^ and previous studies^30^, we observed the highest level of WT *Scn1a* expression in the cortex with expression in cerebellum and hippocampus 34% and 60% lower, respectively. When comparing 1b deletion to WT mice (ANOVA with Tukey’s post-hoc), there was a reduction in *Scn1a* expression as measured by qPCR in 1b^−/−^ mice in all 3 regions tested (reduced by 57% in the cortex, 62% in the hippocampus and 59% in the cerebellum). 1b^+/−^ *Scn1a* expression was not significantly reduced versus WT, but did show a trend towards an intermediate level between WT and 1b^−/−^ levels (**Fig. 2d**).

**Figure 2:**
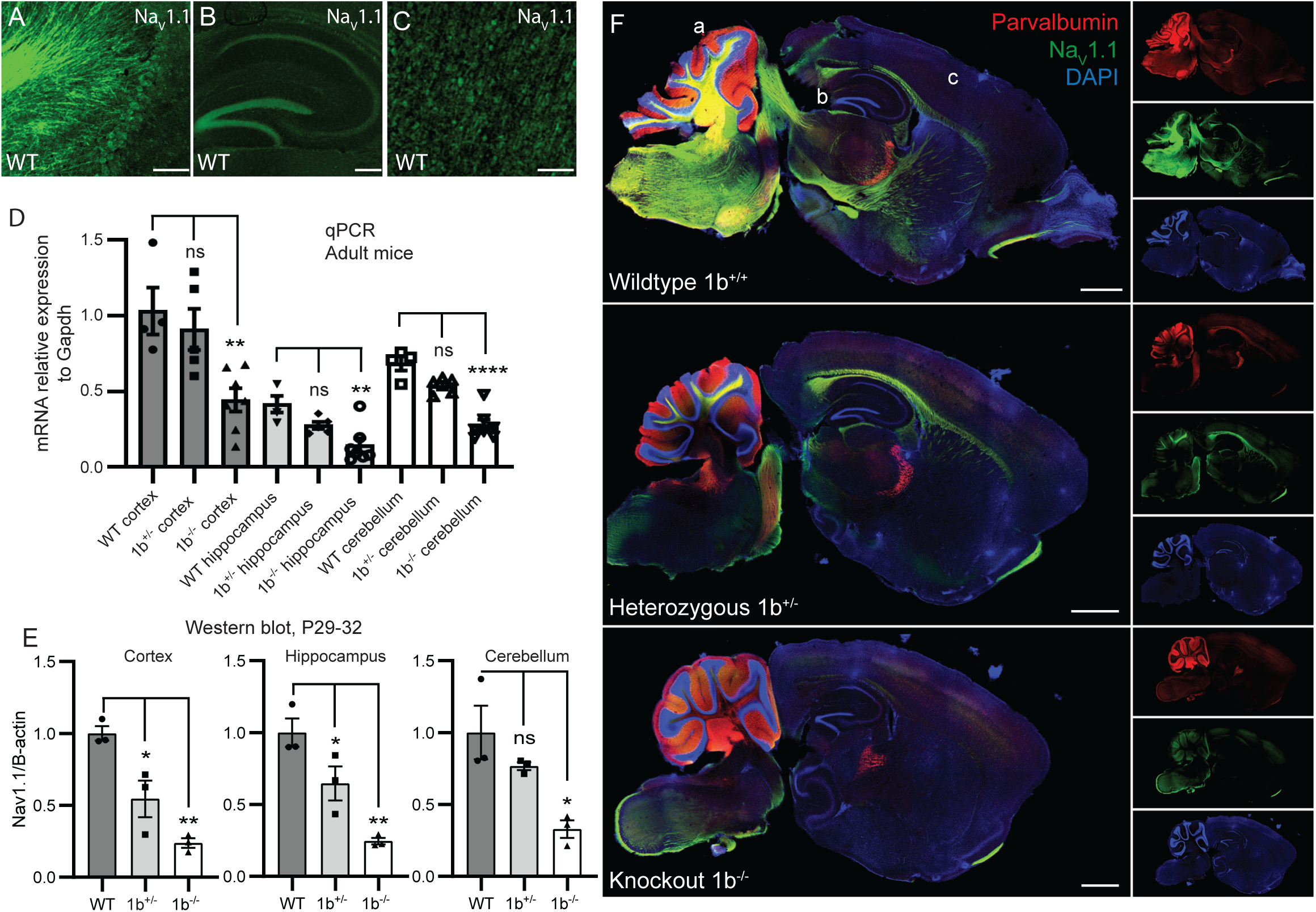
Scn1a expression is reduced in 1b deletion mouse model. (a-c) Immunofluorescent analysis of Na_V_1.1 in wildtype mice across cerebellum (a), hippocampus (b) and cortex (c), regions taken from wildtype in panel f. Scale bars a and b = 100 µm, c = 250 µm. (d) Bar plot showing relative expression of Scn1a using qPCR in 3-month-old mice (mean ± SEM), values normalized to WT cortex. Scn1a expression reduced in 1b-/- cortex vs WT cortex (**P = 0.0092), 1b^−/−^ hippocampus vs WT hippocampus (**P = 0.0029), and 1b-/- cerebellum vs WT cerebellum (****P < 0.0001). (e) Western blots of P29-32 mouse brain membrane fractions, showing reduction of NaV1.1 protein in cortex of 1b+/- (*P=0.0174) and 1b^−/−^ (**P=0.0014) mice, hippocampus of 1b+/- (*P = 0.0445) and 1b^−/−^ (**P = 0.0025) mice and cerebellum of 1b^−/−^ (*P = 0.0142) mice. (f) Immunofluorescent analysis of sagittal sections of P28 mice revealed a reduction in Na_V_1.1 (green) expression in homozygous versus WT mice with no changes in parvalbumin (red) expression. Scale bars = 1 mm.

Reduction in protein expression across the mouse brain was evaluated by Western blot analysis of the membrane fraction from prepared cortex, hippocampus and cerebellum from mice aged P29-32 (n=3 each genotype). 1b^−/−^ mice had significantly decreased Na_V_1.1 protein expression (ANOVA with Tukey’s post-hoc) compared to WT in all three regions, and 1b^+/−^ showed significant decreased expression in cortex and hippocampus and a trend towards reduced levels in cerebellum (**Fig. 2e**). Na_V_1.1 expression was reduced by 45% in 1b^+/−^ mice and 76% in 1b^−/−^ mice in the cortex, 35% in 1b^+/−^ mice and 75% in 1b^−/−^ mice in the hippocampus, and by 23% in 1b^+/−^ mice and 67% in 1b^−/−^ in the cerebellum (**Fig. 2e**). Raw western blots can be seen in supplementary, including blots of the cytoplasmic fraction (**Fig. S2**). These results are consistent with qPCR results, and show that deletion of the extended 1b interval had a larger than expected impact on *Scn1a* and Na_V_1.1 expression considering the proportion of transcripts expected to originate at this element.

With immunofluorescence (IF) we compared qualitative distribution of expression of Na_V_1.1 along with the interneuron marker parvalbumin across 1b^−/−^, 1b^+/−^, and 1b^+/+^ mice at P28 (n=3 each genotype) (**Fig. 2f** and raw images in **Fig. S3**). Notably, deletion of 1b appeared to generally reduce Na_V_1.1 expression, rather than specifically impact certain brain regions, consistent with 5’ RACE TSS activity^30^. Consistent with Western blot quantifications, there was subtle apparent reduction in Na_V_1.1 IF in the brainstem and midbrain between WT and 1b^+/−^ mice, while 1b^−/−^ mice had obvious reduction of expression in the brainstem, midbrain, cerebellum and hippocampus.

### Adult mice harboring heterozygous 1b deletion are susceptible to seizures and exhibit abnormal EEG activity

Given the general failure to thrive, developmental lethality, and spontaneous behavioral seizures in homozygous 1b^−/−^ mice, we focused on heterozygous 1b^+/−^ mice to test for more subtle phenotypes associated with two seizure induction paradigms, exposure to heat and the chemoconvulsant pentylenetetrazole (PTZ), paired with EEG recording. Thermal-evoked seizures are a hallmark of DS and *SCN1A*-associated epilepsy models^45,46^, and thus we tested whether these were induced in 1b^+/−^ mice. P22 1b^+/−^ mice and WT littermates (n=9 per genotype of mixed sex) were subjected to a gradual 0.5 °C increase in body temperature every two minutes from 37.5 °C to 42.5 °C, mimicking the increase in body temperature during a fever. 1b^+/−^ mice displayed heat-induced behavioral seizures at 41.5 °C (**Fig. 3a**) as measured by the Racine scale (**Fig. 3b**) while WT did not present with seizures (**Fig. 3a-b**).

**Figure 3:**
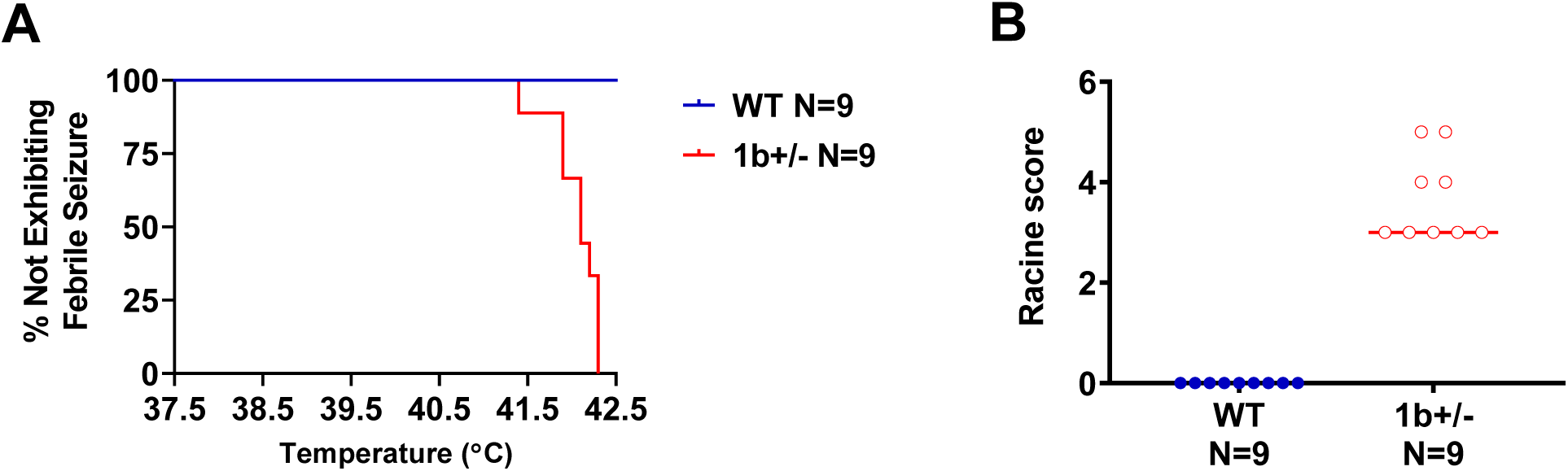
Thermal-evoked febrile seizures in 1b^+/−^ mice and WT littermates. a) The 1b^+/−^ mice began to have seizures at 41.5 °C b) as measured by the Racine scale.

To corroborate and further characterize induced seizure susceptibility, we used a second paradigm, PTZ seizure induction^47^. First, we performed a dose response analysis on mice of the C57BL/6N background strain to identify a PTZ dose that allowed for observations of all stages of behavioral seizure in addition to EEG seizures (**Fig. S4**). To test for induced seizure susceptibility, male and female 1b^+/−^ and WT littermate mice were intraperitoneally administered 80 mg/kg of PTZ. After administration, latencies to first jerk, loss of righting, generalized clonic-tonic seizure, and full tonic extension were measured. In comparison to WT littermates, PTZ-treated 1b^+/−^ mice exhibited increased seizure susceptibility across all measures (**Fig. 4a-d**: first jerk t _(1, 30)_ = 2.171, p = 0.038; loss of righting t _(1, 30)_ = 2.160, p = 0.039; generalized clonic-tonic seizure t _(1, 30)_ = 2.128, p = 0.042; full tonic extension t _(1, 30)_ = 2.207, p = 0.035, unpaired Student’s t-tests).

**Figure 4:**
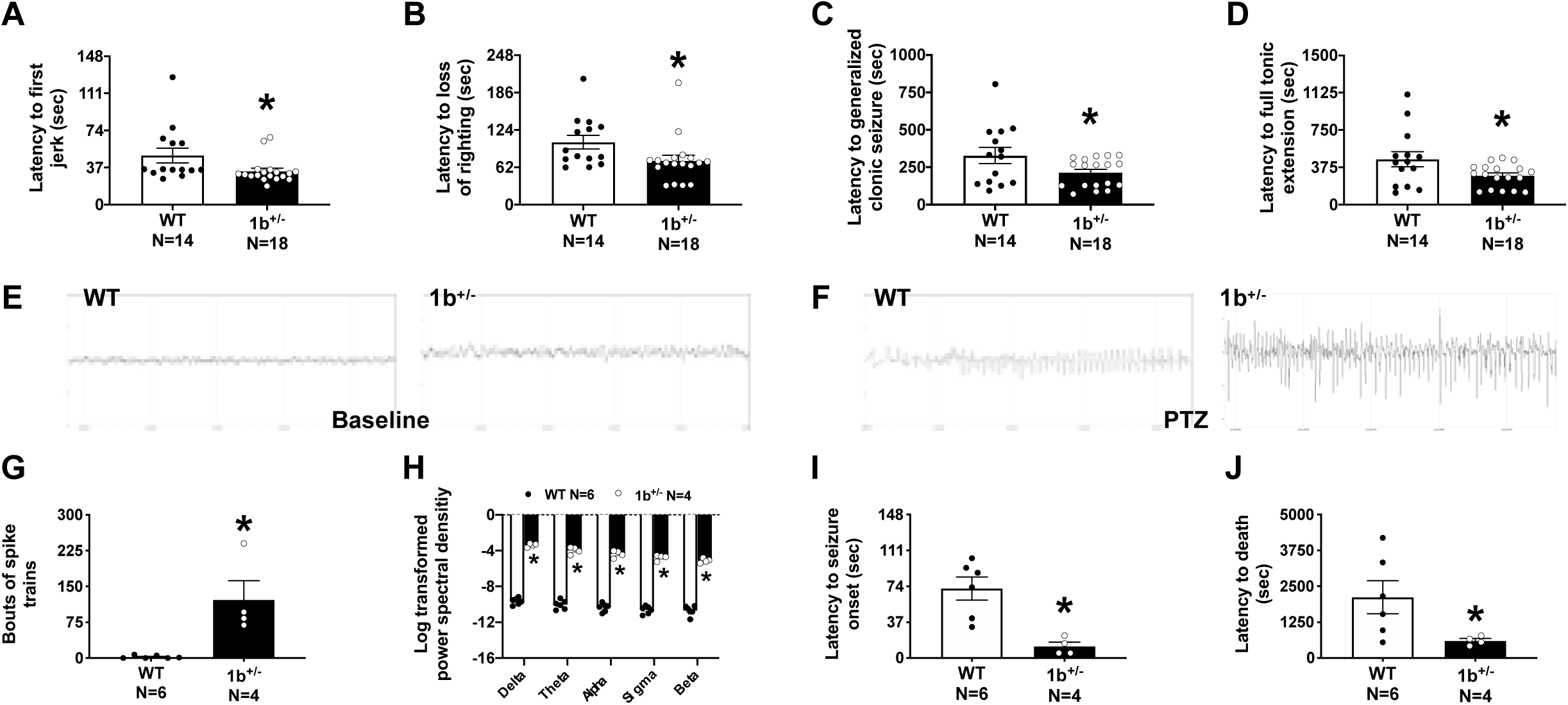
Increased seizure susceptibility and abnormal EEG in heterozygous 1b deletion mice. (a-d) Latency measures were observed after an i.p. injection of 80 mg/kg PTZ over the course of a 30-min trial. Reduced latencies to first jerk, loss of righting, generalized clonic seizure, and full tonic extension were observed in Het mice when compared to WT littermate controls. EEG was collected using a wireless telemetry system before and after an i.p. injection of 80 mg/kg PTZ. (e-f) Representative EEG traces of WT and 1b^+/−^ mice during baseline EEG recording and subsequent PTZ response. Powerband calculations and spiking events were automatically scored. (g) 1b^+/−^ mice had significantly more spiking events and spike trains during baseline EEG acquisition when compared to WT. Scored spiking events are shown on a 1b^+/−^ representative trace and indicated by red lines. (h) 1b^+/−^ mice also had significantly higher power across all frequency bins, Delta (0-4 Hz), Theta (4-8 Hz), Alpha (8-12 Hz), Sigma (12-16 Hz), and Beta (16-30) during baseline when compared to controls. Finally, seizure susceptibility was confirmed with EEG after PTZ administration. (i-j) Reduced latencies to seizure onset and death were observed in 1b^+/−^mice. *, p < 0.05, t-test

Finally, to test for spontaneous neurophysiological phenotypes in 1b^+/−^ mice and to link PTZ-induced behavioral seizures with electrophysiological activity, skull screws for EEG and EMG were implanted in a second group of animals of both sexes and recordings were made over a 24 hour interval, with PTZ induction at the end of the recording period. Comparison of EEG signatures for 1b^+/−^ and WT littermates prior to PTZ administration show elevated spontaneous spiking events and spike trains in 1b^+/−^ mice and increased power spectral density signatures (**Fig. 4e-f**). Spiking activity measured by bouts of spike trains was significantly higher in 1b^+/−^ when compared to WT littermate controls (**Fig. 4g**: t (1, 8) = 3.812, p = 0.005), indicating heightened excitability. 1b^+/−^ deletion subjects also had higher power detected across all frequency bins when compared to WT (**Fig. 4h**: F (1, 8) = 423.9, p < 0.0001, multiple comparisons all had p < 0.0001). At the end of the recording period, PTZ administration in implanted mice reproduced the faster latency to seizure onset and trends towards faster latency to death (**Fig. 4i-j**: t (1, 8) = 3.920, p = 0.004, t (1, 8) = 2.103, p = 0.068, compared to WT by unpaired t-tests), and revealed corresponding increases in EEG activity in response to PTZ. There were no sex differences in induced seizure or EEG outcomes in 1b deletion mice. Overall, these experiments revealed elevated spontaneous EEG spiking and irregular neural signatures, and link PTZ-induced behavioral seizures with increased neurophysiological activity in 1b^+/−^ mice.

### Homozygous but not heterozygous 1b deletion causes cognitive deficits in novel objection recognition (NOR) and spontaneous alternation in the Y-maze

To investigate the impact of 1b deletion on behavior, we performed a tailored battery focused on learning and memory and motor abilities on 1b^+/−^ and surviving 1b^−/−^ male and female mice. 1b^+/−^ mice of both sexes were additionally tested in a comprehensive behavioral battery of standard assays of overall physical health across development, sensorimotor reflexes, motor coordination, anxiety-like, and social behavior to test for subtler phenotypes that might be present in the heterozygous mutants. Details of the administration of behavioral testing is described in the methods, and results from these experiments are summarized in **Table S3**.

Cognitive deficits were observed in 1b^−/−^ but not 1b^+/−^ mice in two corroborating assays of learning and memory, NOR and Y-maze. Following established NOR methods^48,49^ manual scoring by a highly-trained observer blinded to genotype indicated WT and 1b^+/−^ mice spent more time investigating the novel object versus the familiar object, as expected. In contrast, 1b^−/−^ homozygous mice did not exhibit typical novel object preference (**Fig. 5a**: within genotype repeated measures (paired t-test) WT p = 0.002; 1b^+/−^ mice p = 0.018, *, novel versus familiar), illustrating recall of the familiar object and learning. Yet the 1b^−/−^ mice did not exhibit this typical learning and memory (p = 0.8698). Sexes were combined since there was no sex difference observed on time spent sniffing objects (**Table S4**). Control data illustrating no preference for the left or right objects and sufficient time spent investigating the objects is shown in **Fig. 5b**. 1b^−/−^ mice were also impaired on the Y-maze, making less alternation triads compared to WT and 1b^+/−^ mice (**Fig. 5c**: F _(2, 62)_ = 5.693, p < 0.005). Sidak’s multiple comparisons indicated the 1b^−/−^ differed from the WT (p = 0.0192) and 1b^+/−^ (p = 0.0047) mice.

**Figure 5:**
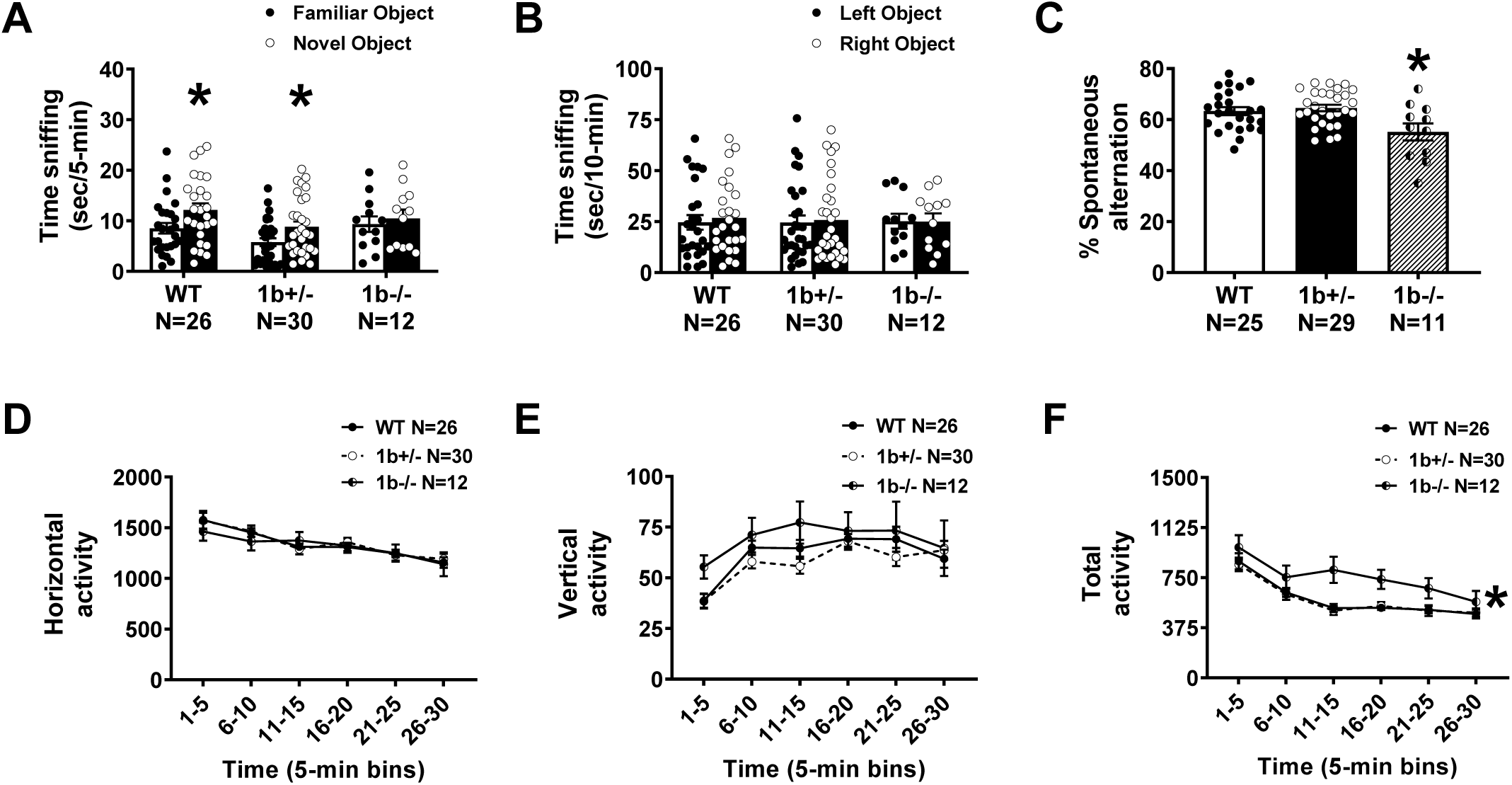
*Scn1a* 1b homozygous deletion mice exhibit learning and memory impairments without confounds in gross motor abilities. Recognition memory was assessed using a novel object recognition assay. (a) 1b^−/−^ mice did not spend more time sniffing the novel object over the familiar object. 1b^+/−^ and WT performed with typical preference. (b) All genotypes showed no preference for either the left or right object during the familiarization phase indicating no innate side bias confounds of lack of object exploration, in the novel object recognition trials. *, p < 0.05, paired-test within genotype using the familiar versus novel object for comparison. (c) Working memory impairments were observed by lower percentages of spontaneous alternation in the Y-Maze. *, p < 0.05, One-way ANOVA. (d) No genotype differences in horizontal (d) or (e) vertical activity counts in the 1b^+/−^ and 1b^−/−^ mice compared to their wildtype littermate controls. (f) 1b^−/−^ deletion mice were hyperactive in total activity during two different 5-min bins of the 30-min assay. Moreover, when total activity is summed and re-graphed as a bar graph, comparisons between 1b^−/−^ versus WT and 1b^+/−^ in total movement were observed. Analyses include both males and females. *p < 0.05, repeated measures ANOVA, main effect of genotype.

Most parameters of gross motor skills and motor coordination were similar across genotypes (**Table S3**). In the open field novel arena assay of locomotion, 1b^−/−^ were hyperactive during 10 minutes of the 30-min session. The time course for horizontal, total and vertical activity was as expected across time, showing normal acclimation to the arena in all genotypes. Horizontal and vertical activity did not differ between genotypes (**Fig. 5d-e**; F _(2, 80)_ = 0.1401, p > 0.05). However, 1b^−/−^ mice were hyperactive in total activity (**Fig. 5f**; F _(2, 80)_ = 5.117, p < 0.008, Two-Way repeated measures ANOVA, genotype x time). Sidak’s multiple comparisons indicated comparisons between the 1b^−/−^ versus the WT mice differed at time of bins 11-15 (p = 0.0014) and differed between the 1b^−/−^ and 1b^+/−^ mice at time bins of 11-15 (p < 0.0001) and 16-20 (p = 0.0270). In addition to highlighting a clear hyperactive phenotype in the 1b^−/−^ mice, linked to DS and numerous NDDs ^50,51^, these results indicate that there were no gross motor abnormalities, inability to rear, or hindlimb weakness that would prevent movement in the learning and memory assays that utilized objects and maze exploration.

1b^+/−^ mice did not exhibit significant consistent phenotypes in a comprehensive battery of assays standard for examining mouse models of NDDs (**Table S3**). For example, 1b^+/−^ mice spent less time in the dark chamber in the light-dark assay (t _(1, 69)_ = 2.121, p = 0.0375, unpaired two-tailed t-test) suggesting elevated anxiety-like behavior, yet no corroborative significance was observed in the plus-maze. Reduced male-female ultrasonic vocalizations in 1b^+/−^ males in the reciprocal social interaction test alludes to aberrant social communication but no corroborative social behavior events were detected (t _(1, 25)_ = 2.143, p = 0.0420, unpaired two-tailed t-test). Also, three chambered approach was typical. Applying rigorous standards used for behavioral studies of NDD models, the absence of two corroborating assays or indices of anxiety-like behavior and sociability precludes us from interpreting these findings as robust phenotypes^52,53^. For all behavioral assays, no sex differences were identified between male and female 1b deletion mice (**Table S4**).

### Differential gene expression in Scn1a 1b deletion mouse hippocampus

We used RNA sequencing (RNA-seq) at P7 and P32 to examine *Scn1a* isoform expression and transcriptional pathology associated with 1b deletion. At P7, forebrain from WT (n=2), 1b^+/−^ (n=4) and 1b^−/−^ (n=2) mice was compared. At P32, analysis was performed on microdissected hippocampus tissue in two rounds. We focused on hippocampus as an example P32 tissue, as hippocampal ablation of *Scn1a* has been specifically linked to seizure and DS-relevant cognitive deficits in mice^54^. For P32, we first compared 1b^−/−^ (n=2) and WT (n=2) mice using 50 bp single end read approach. Next, we compared of WT (n=3) and 1b^+/−^ (n=4) mice using 150 bp paired end read methods in order to more deeply sample reads covering the 1a and 1b UTR region of the *Scn1a* transcript.

At both P7 and P32, *Scn1a* expression showed significant 1b dosage dependent decrease using an additive model (**Table S5-7**). *Scn1a* expression was reduced compared to WT in 1b^+/−^ and 1b^−/−^ mice at P7, though this decrease was only independently significant in 1b^−/−^ mice and failed to pass stringent FDR threshold (P = 0.0047, FDR = 0.28). At P32, where we had increased power due to higher coverage and larger sample numbers and where *Scn1a* expression is much higher in WT, *Scn1a* was significantly lower in both heterozygous and homozygous 1b deletion mice (FDR = 0.00028 and 1.97 x 10^11^, respectively; **Fig. 6a-b, S5a**).

**Figure 6:**
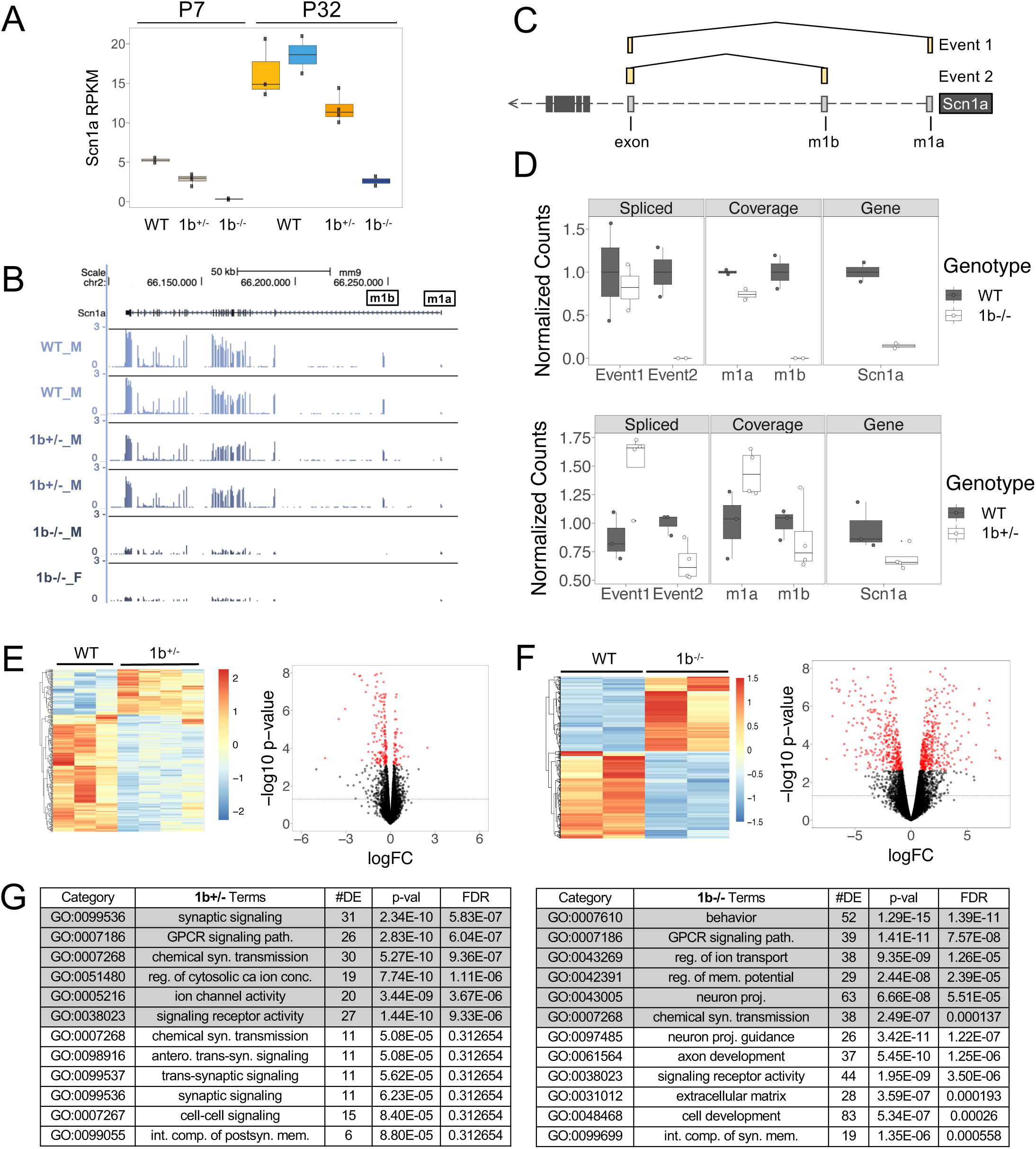
Differential gene expression with *Scn1a* 1b deletion. (a) Bar plot indicating RPKM *Scn1a* expression between WT and 1b^+/−^ or 1b^−/−^ mutants in postnatal day (P) 7 forebrain or P32 hippocampus, (mean ± SEM). (b) Mouse (mm9) *Scn1a* locus showing decrease in coverage in representative P32 heterozygous and homozygous 1b deletion carriers compared to wild-type controls. (c) Schematic showing splicing of m1a and m1b sequences with first *Scn1a* coding exon in reference. (d) Bar plots showing the number of sequencing reads that overlap each splicing event, m1b or m1a locus, and the entire *Scn1a* locus along the x-axis for P32 WT, 1b^+/−^, and 1b^−/−^ mice. The full table is included in the supplement. (e) Heatmap and scatterplot of differentially-expressed genes in P32 1b^+/−^ mice. In the scatterplot, genes with FDR < 0.05 are in red while the dashed line indicates a p-value < 0.05. (f) Heatmap and scatterplot of differentially-expressed genes in P32 1b-/- mice. In the scatterplot, genes with FDR < 0.05 are in red while the dashed line indicates a p-value < 0.05. (g) Table showing select pathways enriched in differentially-expressed genes for P32 1b^+/−^ (left) or 1b^−/−^ (right) mice. Pathways enriched in down-regulated genes are shown in grey. Pathways enriched in up-regulated genes are shown in white. Ontologies are biological pathways (BP), molecular function (MF), or cellular component (CC).

To compare transcripts arising from 1a and 1b at P32, when *Scn1a* expression in WT brain is high, we measured the number of splice junction reads that linked the 1a and 1b non-coding exons with the first *Scn1a* coding exon and the number of total reads that mapped unambiguously to 1a or 1b (**Fig. 6c**). As expected, splice junction and overlapping reads associated with mouse 1b were reduced in 1b^+/−^ mice and abolished in 1b^−/−^ mice (**Fig. 6d, S5b**). Mouse 1a splice junction and overlapping reads appeared reduced in 1b^−/−^ mice, though there were relatively low numbers of reads mapping to these intervals (**Fig. 6d top, S5b left**). In contrast, 1a-associated reads in heterozygous 1b deletion carriers were increased (**Fig. 6d bottom, S5b right**). The increased 1a transcripts present in the 1b^+/−^, but not the 1b^−/−^ mice, is consistent with *trans* compensation by increased expression from the second *Scn1a* allele, but not with compensation of the deleted 1b in *cis* via increased 1a usage. The change in total *Scn1a* reads from RNA-seq at P32 (**Fig. 6d**) was stronger than identified in adult brain tissues via qRT-PCR and Na_V_1.1 western blot. Our findings suggest decreases in RNA and protein levels in 1b deletion mice that are higher than predicted based on the proportion of *Scn1a* transcripts originating at 1b observed here and in previous work^30^.

We tested for differential expression across 15589, 14631, and 15002 genes that were robustly expressed in the P7, P32 heterozygous comparison, and P32 homozygous comparison RNA-seq datasets, respectively (RPKM values in **Table S5-7**). No genes flanking *Scn1a* showed consistent robust differential expression (DE) in 1b^+/−^ and 1b^−/−^ mice at P7 or P32, indicating that the major regulatory effects of the deleted *Scn1a 1b* interval are specific to *Scn1a* expression (**Fig. S5a**). At P7, we identified 21 mostly downregulated DE genes in 1b^+/−^ carriers and 47 downregulated and 61 upregulated DE genes in 1b^−/−^ mice meeting an FDR < 0.05 threshold (**Table S8, S9**). The small effect of 1b deletion on differential gene expression at P7 is consistent with the low expression and non-essential role of *Scn1a* in early postnatal development^55^. Later in development, P32 heterozygous *1b* deletion was associated with 223 DE genes (175 downregulated, 48 upregulated) at FDR < 0.05 (**Fig. 6e, Table S10**). Homozygous *1b* deletion carriers exhibited much stronger transcriptional impact, with a total of 723 DE genes (337 downregulated, 386 upregulated) DE at FDR < 0.05 (**Fig. 6f, Table S11**). Volcano plots that show log2 fold change effect sizes and significance values for DE genes in 1b deletion mice shown in **Fig. 6e and 6f**. Gene set enrichment analysis of Gene Ontology (GO) found general synaptic signaling and function were enriched among downregulated DE genes in P32 heterozygous carriers, with no terms passing FDR < 0.05 criteria for upregulated genes. In P32 homozygous *1b* carriers, enriched terms for neuron development and differentiation were associated with upregulated DE genes while synaptic signaling and mature neuronal function terms were enriched among downregulated DE genes (**Fig. 6g**). Heterozygous and homozygous *1b* deletion mutants shared 104 DE genes (**Fig. S5c**), which were primarily downregulated and enriched for synaptic and differentiation terms (**Fig. S5d**).

## Discussion

The majority of focus and functional studies of alternative promoters have been on genes where the multiple alternative TSSs are predicted to have discrete cell-type or tissue-specific activity^6–9^. However, recent studies of TSS usage and promoter interactions suggest a model where alternative promoters interact physically and are co-active in the same cells^12–14^. In these situations, it is largely unknown what the requirement for individual TSS and associated regulatory DNA may be. Here we focus on one specific putative non-canonical disease-relevant alternative promoter, a 3-kb evolutionarily-conserved DNA region including the previously described *Scn1a 1b* TSS. We show that full deletion of this interval from the mouse genome causes significant decrease in *Scn1a* expression and Na_V_1.1 protein, and results in spontaneous seizures and high developmental lethality with significant cognitive and behavioral deficits in surviving mice. While phenotypes were less severe in heterozygous 1b^+/−^ mice, presence of temperature and PTZ-induced behavioral seizures and elevated observations of epileptiform indices in EEG in these mice indicate a milder phenotype with relevance to SCN1A-associated epilepsies^56–58^. Our results define 1b as an essential disease-relevant *Scn1a* regulatory region and show that loss of regulatory DNA associated with a non-canonical TSS can have a surprisingly strong and translationally-relevant phenotypic impact.

There are multiple possible explanations for the observed strong impact of loss of the 1b interval on *Scn1a* expression. First, 1a and 1b isoforms may indeed be discretely regulated, but previous measures of 1b-originating transcripts must have significantly underestimated the actual contribution of 1b transcripts to *Scn1a* expression. However, there is no evidence that earlier studies were incorrect and our estimates of *Scn1a* overall and 1a and 1b RNA-seq read frequency in 1b deletion mice do not support this simple model. Alternatively, the 1b TSS could have increased activity earlier in development or 1b-associated regulatory DNA activity could also be required for 1a transcription. These models are both plausible and consistent with our results. Considering the frequency of promoter-promoter interaction and reported common co-expression of alterative TSSs in single neurons, many brain genes could share similar regulatory structure as *Scn1a*. Our findings are consistent with models suggesting regulatory DNA at putative alternative promoters and associated regulatory sequences contributes to transcriptional activation across interacting TSSs. Further experiments are needed to resolve the specific cellular, molecular, and developmental function and potential co-dependence of the 1a and 1b intervals and associated transcripts, and studies of other genes are needed to test if this phenomenon is widespread. Our findings represent initial insights into the essential regulatory roles of non-canonical promoters even when such TSSs produce mRNA encoding identical amino acid products.

Annotation of the genome has led to major gains in understanding transcriptional wiring, yet it has been surprisingly difficult to predict the sufficiency and necessity of specific regulatory elements, even those expected to be critical based on comparative and functional genomics^3,10,59^. Knockout mouse models have been a gold standard for testing the phenotypic consequences of mutations, and recent efforts deleting non-coding DNA have provided critical insights into the role of regulatory DNA^59–62^. Here, we used CRISPR/Cas9-mediated deletion to assess the role of the evolutionarily-conserved 1b interval on higher order neurological phenotypes in mice. Homozygous 1b deletion caused spontaneous seizures and behavioral deficits and had a strong impact on survival, demonstrating the essential nature of the deleted interval. Further studies are needed to define the minimal and core nucleotides within the 1b interval and to define proteins that bind and participate in regulation. In addition, similar functional studies of other *Scn1a* regulatory DNA elements, and specifically of the 1a region, are necessary to determine which regulatory DNA regions are necessary and sufficient for expression in the brain.

Heterozygous loss of the 1b interval appears to have a less severe impact compared to truncating *Scn1a* mutations, which are sufficient to reduce survival and cause behavioral and cognitive deficits relevant to DS and NDD in mice^20,26^. It is possible that phenotypes are milder in heterozygous 1b deletion mice in this study compared to other *Scn1a* mouse models due to differences in genetic background or environment. In a review of *Scn1a*^+/−^ mouse models of Dravet syndrome^63^ it was noted that those bred on a 6N background were more susceptible to hyperthermia-induced seizures, yet had milder spontaneous seizures and improved survival rates relative to 6J crosses. DS-model mice bred on 129/SvJ genetic background have a higher threshold for thermally-induced seizures, no cognitive impairments and reduced rates of premature death^64^. The 1b deletion mouse line was generated on a C57BL/6N background and thus it is possible that phenotypes are milder due to this compared to if they were bred with C57BL/6J. However, there is also evidence that the milder phenotypes in 1b deletion mice relative to *Scn1a* coding loss-of-function mutants is due to the different impacts on *Scn1a* dosage. Previously reported phenotypes of mice harboring heterozygous DS-associated *Scn1a* truncating mutations are similar to or less severe than homozygous 1b deletion phenotype identified here, suggesting stronger phenotypic impact in line with haploinsufficiency is produced by the more severe reduction in *Scn1a* expression caused by homozygous 1b deletion. In support of this, the P32 homozygous DE signatures of downregulated synaptic expression and upregulated expression of earlier neuronal differentiation and maturation genes are consistent with previous data on *Scn1a* truncating mutants^65^. Notably, these DE results could be driven by either developmental changes or reflect seizure pathology in 1b deletion mice^65^. Further studies of 1b deletion and *Scn1a* coding mutant mouse lines on the same genetic background will be needed to directly compare phenotype severity associated with 1b deletion.

While we did not identify spontaneous behavioral seizures or corroborated behavioral phenotypes in heterozygous 1b deletion mice, our characterization of 1b^+/−^ mice showed general epilepsy relevance. Heterozygous 1b deletion mice exhibited spontaneous spike trains and abnormal EEG spectral bandwidths. In particular, the EEG results with increased spike trains observed 1b^+/−^ mice are indicative of general seizure relevance. It is likely that the increased spike events at baseline are associated with susceptibility to thermally-induced seizures seen in 1b^+/−^ mice. At 41.5 °C, the temperature of seizure onset is higher than those observed for global Scn1a^+/−^ mice (38.5 ± 0.2 °C) ^66^ and hippocampal Na_V_1.1 deletion mice (40.3 ± 0.2 °C)^54^. There are neurodevelopmental disorder genetic models where spontaneous behavioral seizures are not seen in mice while they are in humans^67–69^. As female 1b^+/−^ failed to efficiently reproduce, it is possible that milder but still DS-relevant behavioral and cognitive deficits are present in heterozygous 1b deletion mice. The degree to which 1b deletion is directly relevant to DS will require further studies. Regardless, our findings show the relevance of this novel *Scn1a* regulatory deletion mouse line to *SCN1A*-associated epilepsy.

The EEG spectral phenotypes in heterozygous 1b deletion mice overlap with other neurodevelopmental disorder and epilepsy models, and represent a potential translational biomarker for future investigation. Elevated delta spectral power is a biomarker of Angelman Syndrome (AS)^70^ and elevated beta spectral power is posited to be a biomarker of Dup15q syndrome^71,72^. These disorders are of interest as there are co-occurring features with DS and epilepsy. AS and Dup15q both have high rates of seizures, cognitive disruption and comorbid diagnosis with autism. Neural signatures in EEG by power bands can be similarly measured in both rodents and humans, and thus our findings have translational relevance^56–58^. Analysis of spike-firing and oscillatory activity during rewarded trials in touchscreen assays have recently been described in detail^73^. Given the behavioral deficits in cognitive function and firing activity identified here, future studies investigating behavioral outcomes and neurophysiological signals are warranted and will shed light on relationship between 1b deletion, EEG spectral phenotypes, and behavior.

*SCN1A*-associated epilepsies, including DS, remain difficult to treat as conventional sodium channel blockers are usually ineffective and may even exacerbate the disease^29,74^. Precision therapies that rescue Na_V_1.1 haploinsufficiency in relevant cell types would be preferred to ameliorate symptoms and reduce side effects compared to more globally acting therapies. Using CRISPR/dCas9 induction, we increased *SCN1A* expression in HEK293 cells by targeting the 1b region. Application of a similar synthetic transcriptional activation therapeutic strategy has shown exciting promise in vivo in mice, where a dCas9-based activator combined with locus-specific guide RNA delivered to hypothalamus was capable of rescuing obesity phenotypes in *Sim1* and *Mc4r* heterozygous mutant mice^75^. dCas9-VP160 activation of *Scn1a* 1b, but not 1a, by a single sgRNA was able to enhance *Scn1a* expression in P19 cells^76^. Delivery of the guide and dCas9-VP64 via AAV intracerebroventricular injections to forebrain GABAergic interneurons in a model of Dravet syndrome ameliorated hyperthermia-induced seizures^76^. Our data demonstrates that 1b is an essential and important regulator of *Scn1a* expression and highlights a potential target for epigenomic intervention in *SCN1A*-related epilepsies. Studies characterizing the regulatory DNA at disease-relevant loci, as we have done here with the *Scn1a* 1b region, will be required to properly design therapies using targeted expression rescue.

The work here on the *Scn1a 1b* regulatory region contributes to functional dissection of the regulatory wiring of a major epilepsy risk gene. Our findings show that *Scn1a* 1b regulatory deletion mice represent a general epilepsy-relevant model that will be valuable for understanding the relationship between *Scn1a* dosage and neurological phenotypes in a genetic preclinical model. Our study justifies increased focus on non-coding regulatory DNA in genetic screening of DS and epilepsy patients, and highlights the need for more in-depth functional studies of regulatory DNA elements in general and specifically in haploinsufficiency associated disorders.

## Acknowledgments

Sequencing was performed at the UC Berkeley and UC Davis DNA cores. This work was supported by funds from the Dravet Syndrome Foundation, Simons Foundation Autism Research Initiative and by NIGMS R35 GM119831. L.S.-F. was supported by the UC Davis Floyd and Mary Schwall Fellowship in Medical Research and by grant number T32-GM008799 from NIGMS-NIH. A.A.W. was supported by Training Grant number T32-GM007377 from NIH-NIGMS and F31 MH119789-01. R.C.-P. was supported by a Science Without Borders Fellowship from CNPq (Brazil). A.V., L.A.P. and D.E.D. were supported by National Institutes of Health grants R24HL123879, U01DE024427, R01HG003988, U54HG006997 and UM1HL098166. Research conducted at the E.O. Lawrence Berkeley National Laboratory was performed under Department of Energy Contract DE-AC02-05CH11231, University of California. We also thank Heather Boyle at the MIND Institute for her diligence and assistance with the mouse colonies. This work was supported by generous funding from the NIH R01NS097808 (JLS, NAC, AA) and the MIND Institute’s Intellectual and Developmental Disabilities Resource Center (HD079125, PI Abbeduto).

## Author contributions

JLH and AA are listed as joint first authors, as each led major components of the study. JLH, AA, NAC, TS, AAW, RCP, LSF, JC, JLS and ASN designed the experiments and analyses. JLH, AA, NAC, TS, IZ, SM, TAF, AN, DQ, MS, EK, SA, JC, AG, JTL, CPC, LAP, AV and DED performed experiments. AA, NAC and TAF carried out mouse behavior. LSF, AAW, ASN performed and interpreted transcriptomic analysis. JLH, IZ, SM, AN, DQ, SA, MS, JC ran cell culture experiments. JLH, AA, NAC, JLS and ASN drafted the manuscript. All authors contributed to the revisions.

---

**Figure S1:**
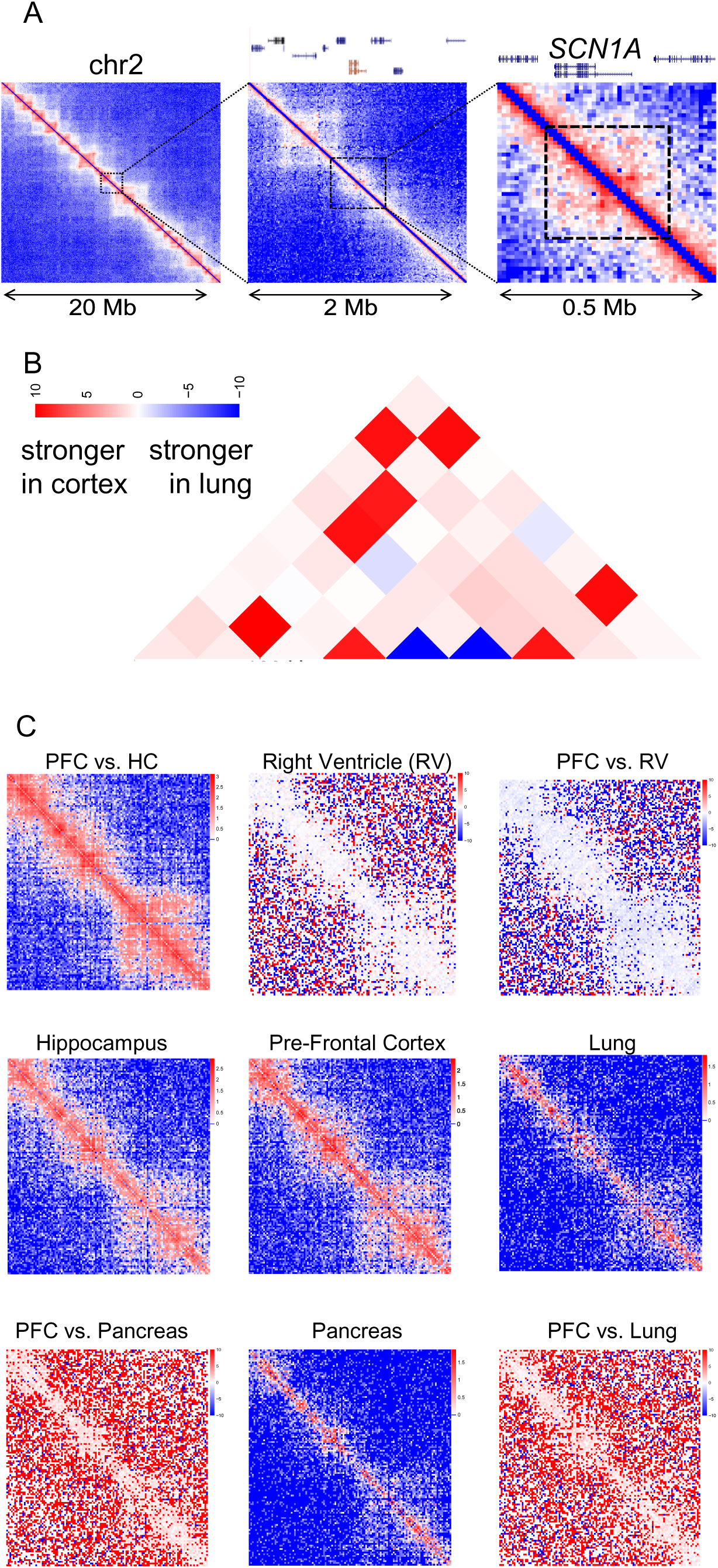
Tissue and brain regional differences in chromatin conformation. (a) Hi-C contact heatmaps showing chromosomal neighborhood of *SCN1A* at different ranges. (b) Contrasting differential Hi-C contact heatmaps showing differences between PFC and lung. (c) Hi-C contact map heatmaps at 40-kb resolution for a 5-Mb region around *SCN1A* gene. In the central rows and columns are the absolute contact maps, while the corner plots represent the differential contact maps between pre-frontal cortex (PFC) and the indicated tissue. For the latter, red color indicates stronger contacts in PFC in relation to the other tissue, and blue color the opposite.

**Figure S2:**
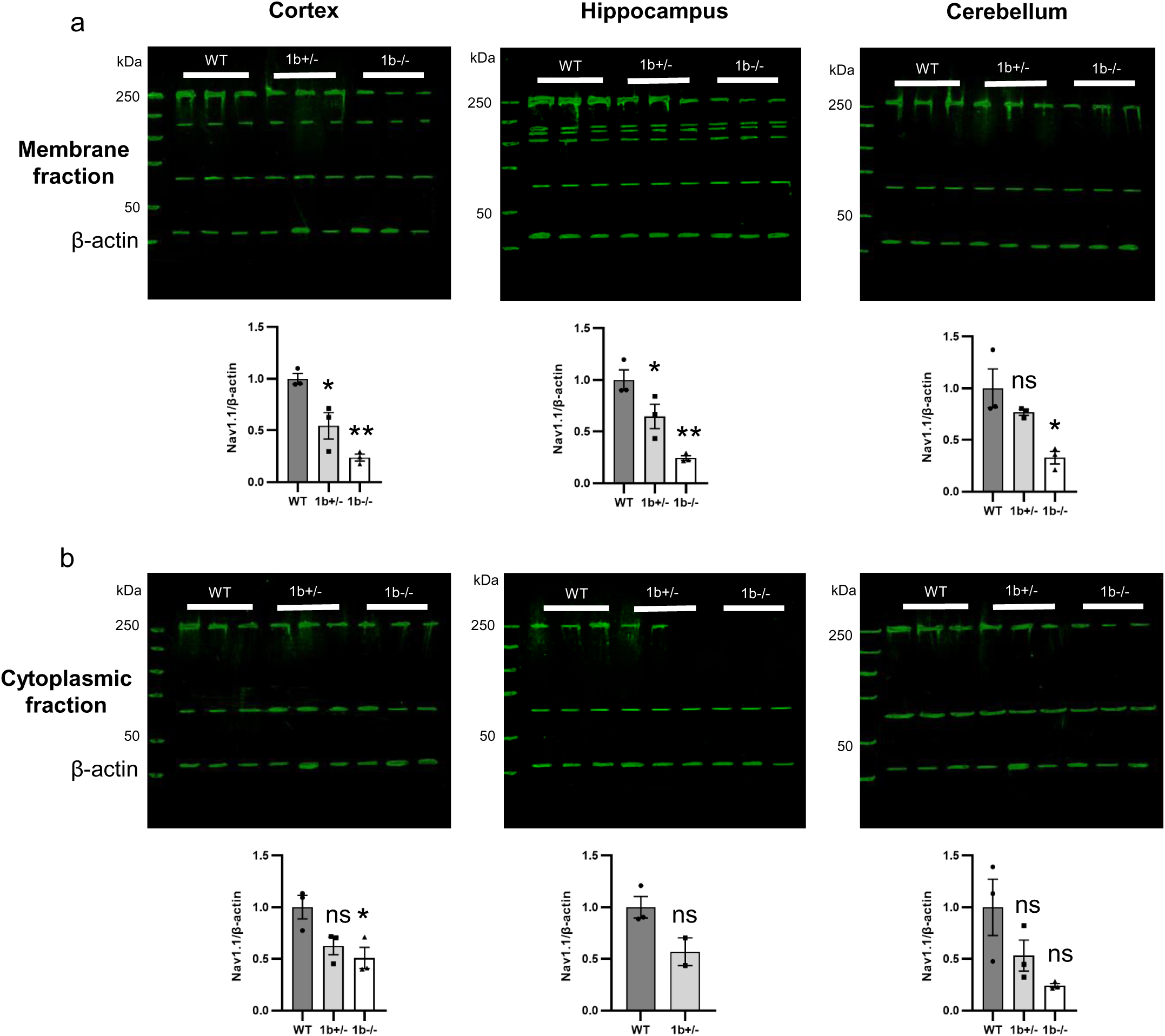
Western blots of Na_V_1.1 (250 kDa) and β-actin (45 kDa) proteins in P29-32 brain lysates, membrane and cytoplasmic fractions ran separately. a) Membrane fraction western blot for cortex shows decrease in Na_V_1.1 protein abundance in 1b^+/−^ (*P=0.0174) and 1b^−/−^ (**P=0.0014) mice. In the hippocampus there is also a decrease in 1b^+/−^ (*P=0.0445) and 1b^−/−^ (**P=0.0025) mice. In the cerebellum there was a decrease in Na_V_1.1 in 1b^−/−^ (*P=0.0142) compared to WT. b) Cytoplasmic fraction western blot shows a decrease in Na_V_1.1 in 1b^−/−^ (*P=0.0321) versus wildtype, no other significant changes. WT n=3, 1b^+/−^ n=3, 1b^−/−^ n=3. ANOVA with Tukey’s post-hoc, significance is versus WT; cytoplasmic hippocampus uses unpaired t-test due to no bands in the KO samples. Error bars represent mean ± SEM. Bands not at 250 and 45 kDa are non-specific and do no show significant changes between genotypes.

**Figure S3:**
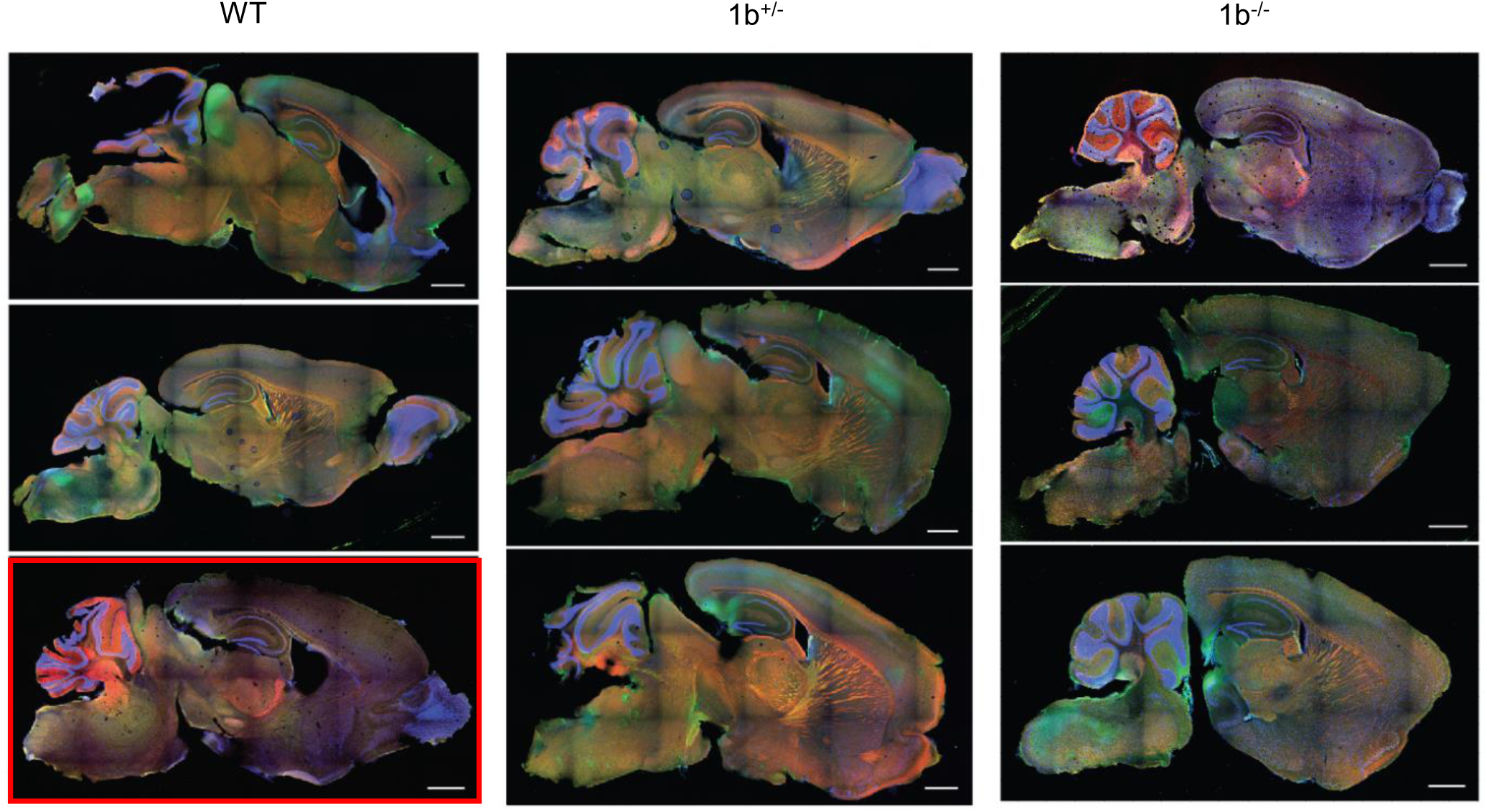
Immunofluorescent analysis of sagittal sections of P28 mice. WT, heterozygous (1b^+/−^) and knockout (1b^−/−^) 1b mice (n=3 each genotype) underwent immunohistochemistry (IHC) for Na_V_1.1 (green) and parvalbumin (red) expression (DAPI shown in blue). No observable difference between WT and 1b^+/−^. In 1b^−/−^ mice Na_V_1.1 appears reduced in brainstem (2/3 KO), corpus callosum (3/3) and hippocampus (3/3). No observable difference at the IF level in the cortex. WT in red box presented patchy labelling. Scale bars = 1 mm.

**Figure S4:**
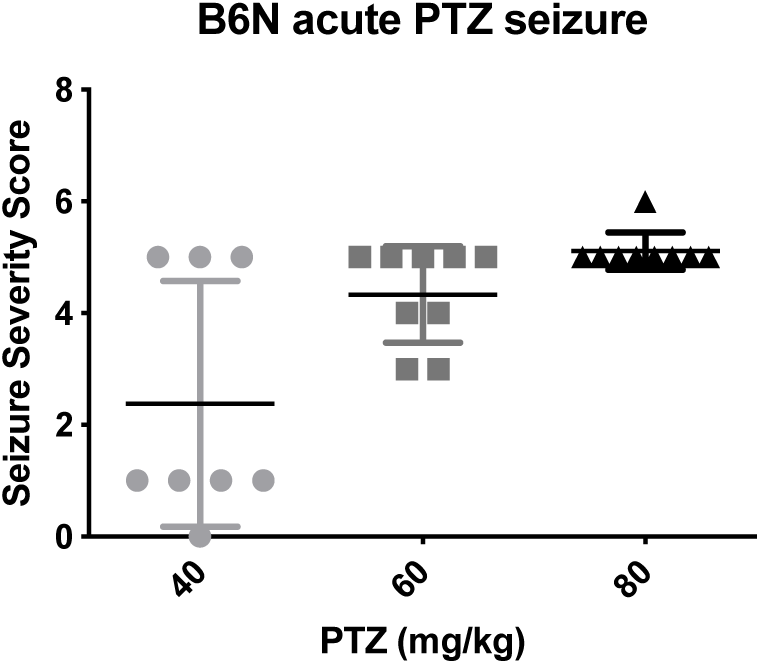
Seizure severity following Pentylenetetrazole (PTZ) administration in C57BL6/N mice. 80 mg/kg was chosen to ensure all seizure stages were reached.

**Figure S5:**
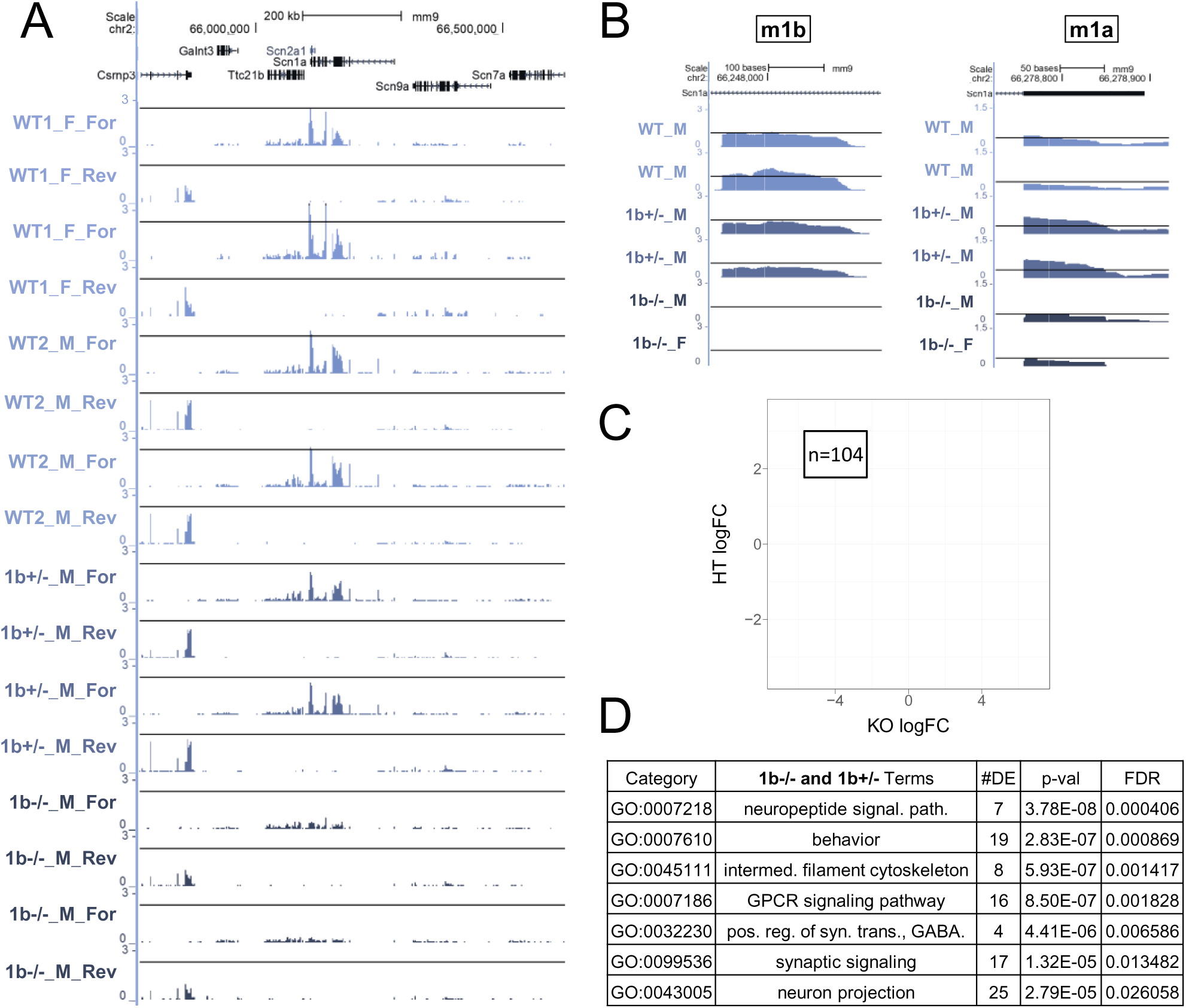
Differential gene expression with *Scn1a 1b* deletion in P32 hippocampus. (a) Mouse (mm9) *Scn1a* and nearby-gene loci for P32 wild-type, 1b^+/−^, and 1b^−/−^ mice. WT1 refers to mice sequenced with the 1b^−/−^ mutants. WT2 refers to mice sequenced with 1b^+/−^ mutants. (b) Coverage of the m1b (left) and m1a (right) loci across representative P32 wild-type and mutant mice. (c) Scatterplot of differentially-expressed genes meeting a p-value < 0.05 and fold-change > 1.5 threshold in both P32 heterozygous and homozygous 1b deletion datasets. (d) Table showing select pathways enriched in genes having shared differential-expression in both P32 mutant mouse datasets. Ontologies are biological pathways (BP), molecular function (MF), or cellular component (CC).

## Materials and Methods

### Generation of 1b mutant mice

We used Cas9-mediated mutagenesis of C56BL/6N oocytes to generate a mouse line harboring deletion of a conserved portion on the noncoding region of *Scn1a* containing the previously described 1b^30^ regulatory region. Guide RNA was designed and synthesized according to described methods^63^, pooled with Cas9 mRNA and injected into mouse oocytes. The gRNA sequences were GGAGATCTGGGTAGTCCTCG and GCTTTTCATACTATAGTGAG. Initial Cas9 targeting was performed at Lawrence Berkeley National Laboratory. F0s (induced on C57BL/6N background) carrying mutations were genotyped and bred to expand a line harboring a 3063 bp deletion at the 1b interval (mm10 - chr2:66407567-66410630).

The colony was rederived and maintained by crossing male 1b deletion carriers with C57BL/6N wild-type females (Charles River). Extensive crossing of heterozygous mutation carriers to wild-type animals vastly reduces the likelihood that any potential off-target mutations caused by Cas9 targeting would persist in our 1b deletion line. Genotypes were identified via PCR and sequence-verified for all animals included in analyses, with the primers in Table 1. All mouse studies were approved by the Institutional Animal Care and Use Committees at the University of California Davis and the Lawrence Berkeley National Laboratory. Subject mice were housed in a temperature-controlled vivarium maintained on a 12-h light–dark cycle. Efforts were made to minimize pain and distress and the number of animals used. Survival was monitored and log-rank Mantel Cox used to assess survival rate.

**Table 1:**
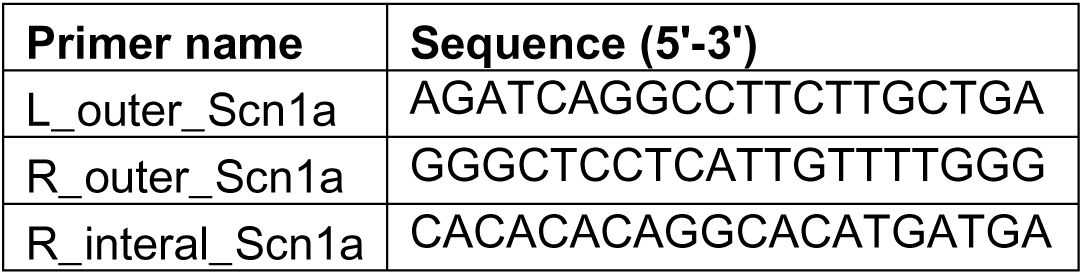
Scn1a 1b deletion genotyping primers.

**Table 2.** below provides the sex, genotype and n for each experiment.

| Experiment | n | Region | Age |
| --- | --- | --- | --- |
| qPCR | WT=4F, 1b+/-=5<br>(4F,1M), 1b-/-=7<br>(2F,5M) | Cortex<br>Hippocampus<br>Cerebellum | 3<br>months |
| WB | WT=3M, 1b+/-=3M,<br>1b-/-=3M | Cortex<br>Hippocampus<br>Cerebellum | P29-P32 |
| IF | WT=3, 1b+/-=3, 1b-/-<br>=3 | Brain | P28 |
| RNA-seq | WT=2M, 1b+/-=4 (2M,<br>2F), 1b-/-=2F | Forebrain | P7 |
|  | WT=2F, 1b-/-=2 (1M,<br>1F); WT=3M, 1b+/-=4<br>(3M, 1F) | Hippocampus | P32 |
| Behavior | WT=26(12M, 14F),<br>1b+/-=30(15M, 15F),<br>1b-/-=12 (6M, 6F) |  | Started<br>at 6<br>weeks |
| Thermal-<br>induced<br>seizures | WT=9(5M, 4F)<br>1b+/-=9 (6M, 3F) |  | P22 |

### RNA collection

Cortex, hippocampus and cerebellum were regionally dissected from one hemisphere of P7, P32 and 3-month-old homozygous deletion, heterozygous and wildtype mice. Both male and female mice were used, though there was not equal sex representation across genotypes. Total RNA was isolated using RNAqueous kit (Ambion) and assayed using an Agilent BioAnalyzer instrument.

### qRT-PCR

Differential expression of *Scn1a* was verified by qRT-PCR at 3 months old. Primers are reported in Table 3 and qPCR was performed with SYBR green PCR master mix (Applied Biosystems). Samples were excluded if technical replicates failed. Cycle counts were normalized to Gapdh. Statistical analysis was performed using ANOVA followed by Tukey’s on relative gene expression between genotypes using ΔΔCT.

**Table 3:**
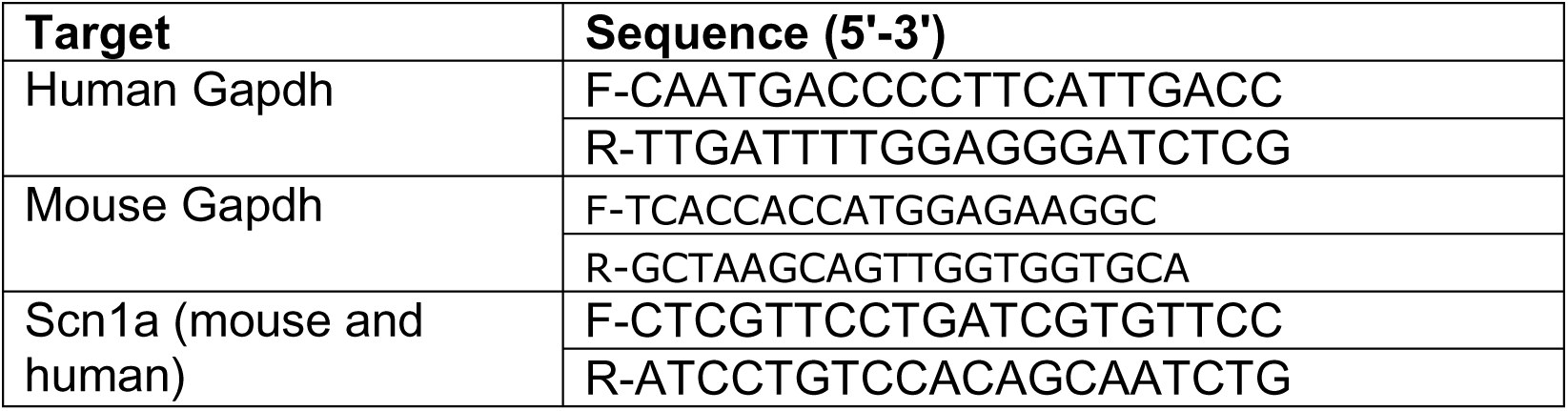
qPCR primers.

RNA from P7 forebrain and P32 hippocampus was collected as described above. Samples included males and females of each genotype when possible. Stranded mRNA sequencing libraries were prepared using TruSeq Stranded mRNA kits. Each round of sequencing was, quantified, pooled, and sequenced in one lane on the Illumina HiSeq platform using a single-end 50-bp (P7, P32 homozygous) or paired-end 150 (P32 heterozygous) strategy at the UC Davis DNA Technologies Core.

The transcriptomic analysis was performed as before^64^. Reads from RNA-seq were aligned to the mouse genome (mm9) using STAR (version 2.7.2)^65^. Aligned reads mapping to genes were counted at the gene level using subreads featureCounts^66^. The mm9 knownGene annotation track and aligned reads were used to generate quality control information using the full RSeQC tool suite^67^. Unaligned reads were quality checked using FastQC.

### Differential expression analysis

Raw count data for all samples were used for differential expression analysis using edgeR^68^. Genes with at least 1 read per million in at least one sample were included for analysis, resulting in a final set of 15589, 14631, and 15002 genes for differential testing in P7 and P32 mice, respectively. Multidimensional scaling analysis indicated that *Scn1a* expression and sex were the strongest driver of variance across samples. Tagwise dispersion estimates were generated and differential expression analysis was performed using a generalized linear model with genotype as the variable for testing. Effect of genotype was modelled as individual comparison of heterozygous and homozygous 1b deletion mice with the respective WT controls. Normalized expression levels were generated using the edgeR pseudocount and rpkm functions. Aligned reads contained in BAM files from each sample were counted to calculate the overlap of sequencing reads with each locus. The coordinates for each locus were m1a: chr2:66,278,753-66,278,887, the m1b deletion region: chr2:66245632-66248697, and m1c: chr2:66,249,400-66,249,514. Mouse gene ontology (GO) data was downloaded from Bioconductor (org.Mm.eg.db). We used the goseq package to test for enrichment of GO terms indicating parent:child relationships. For GO analysis, we examined down- and up-regulated genes separately for genes meeting an FDR < 0.05. For the enrichment analysis, the test set of differentially expressed genes was compared against the background set of genes expressed in our study.

### Immunofluorescence

All histological experiments were performed at least in triplicate and experimenters were blinded to genotype. Following anesthesia, P28 male and female mice were transcardially perfused with 4% paraformaldehyde (PFA) in HEPES, followed by overnight fixation in the same solution. After fixation, brains were removed from the skull, embedded in 2% LTE agarose/ Tris-buffered saline (TBS) and cut coronally in 50 µm sections on a vibratome (VT 1000S, Leica). Sections underwent antigen retrieval in a solution of 0.1 M sodium citrate (pH 6), 200 mM sucrose and 1% (v/v) hydrogen peroxide at 60⁰C for one hour. Subsequently, sections were permeabilized and blocked in TBS with 0.1% Triton X-100 and 5% normal donkey serum for 24 hours at room temperature. Immunolabelling was carried out using primary antibodies directed against Na_V_1.1 (K74/71, mouse, IgG1, 1:100, NeuroMab) and parvalbumin (L114/3, 75-455, mouse, IgG2a, 1:100, NeuroMab). Subclass specific secondary antibodies (488 and RRX) were used at 1:200 (Jackson ImmunoResearch Laboratories Inc.). All imaging was carried out on a Nikon A1 laser scanning confocal microscope. FIJI (National Institutes of Health) was used for image processing with settings consistently applied to across samples, raw tile scans in Supplementary Fig. 3.

### Western blot

Flash frozen samples (n=3 per genotype) were prepared for Western blot using the Mem-PER Plus protein extraction kit (89842, Thermo Scientific) to isolate membrane and cytoplasmic protein fractions from mice aged P29-32. We ran 40 µg of protein on 10% gels using the Mini-PROTEAN Tetra Cell western blotting system (Bio-Rad). Anti-Na_V_1.1 (Ab5204a, rabbit, 1:1000, Millipore) and anti-beta-actin (Ab8227, rabbit, 1:5000, Abcam) primary antibodies were incubated overnight in Odyssey blocking buffer (LI-COR). Secondary antibodies (IRDye 800CW Donkey anti-Rabbit IgG Secondary Antibody) were used at 1:5000 in Odyssey blocking buffer (LI-COR) for one hour at room temperature. Blots were visualized using a LI-COR Odyssey CLx system and quantified in FIJI. Protein levels assayed via western blot were compared by one-way ANOVA and Tukey’s post-hoc.

### Mouse colony at UC Davis Medical Center

Heterozygous (+/-) breeders were transferred from the UC Davis Center of Neuroscience to the UC Davis Medical Center. Offspring were maintained on the C57BL/6N background from The Jackson Laboratory (Bar Harbor, ME). Colonies were maintained with two breeding paradigms: wildtype (+/+) by heterozygous (+/-) and heterozygous (+/-) by heterozygous (+/-) crosses, giving rise to wildtype (+/+), heterozygous (+/-), and knockout mice (-/-). After weaning on PND 21, mice were socially housed in groups of 2-4 by sex. Cages were housed in ventilated racks in a temperature (68-72°F) and humidity (∼25%) controlled vivarium on a 12:12 light/dark cycle with lights on at 07:00, off at 19:00-h. Standard rodent chow and tap water were available ad libitum. In addition to standard bedding, a Nestlet square, shredded brown paper, and a cardboard tube were provided. All subjects were tested between 2-5 months of age. All measures were conducted by an experimenter blind to genotype.

### Behavioral Assay Design

Both male and female subjects were used in this study. Subjects (WT – Males N=12, Females N=14, HET – Males N=15, Females N=15, and KO – Males N=6, Females N=6) began the behavioral battery at 6-week of age. All behavioral tests were performed between 09:00 and 17:00-h during the light phase of the 12:12 light/dark cycle. Mice were brought to an empty holding room adjacent to the testing area at least 1-h prior to the start of behavioral testing. To minimize the carry-over effects from repeated testing, assays were performed in the order of least to the most stressful tasks. Subjects were sampled from HET x HET and HET x WT pairings. The order and age of testing was as follows with at least 48-hours separating tasks: (1) Elevated plus maze at 6 weeks of age, (2) light dark conflict at 6 weeks of age, (3) open field at 7 weeks of age, (4) beam walking at 7 weeks of age, (5) rotarod at 8 weeks of age, (6) novel object recognition at 9 weeks of age, (7) spontaneous alternation at 10 weeks of age, (8) self-grooming at 11 weeks of age, (9) social approach at 11 weeks of age, (10) male-female social interaction at 12 weeks of age, (11) acoustic startle at 13 weeks of age, (12) pre-pulse inhibition at 13 weeks of age, and (13) fear conditioning at 14 weeks of age..

### Developmental milestones

Developmental milestones were measured on PND 2, 4, 6, 8, 10, and 12, as previously^46,69^. Body weight, length (nose to edge of tail), and head width were measured using a scale (grams) and a digital caliper (cm). Cliff avoidance was tested by placing each pup near the edge of a cardboard box, gently nudging it towards the edge, and measuring the time for it to turn and back away from the edge. Failures to avoid the cliff was recorded as a maximum score of 30-s. Righting reflex was tested by placing each pup on its back, releasing it, and measuring the time for it to fully flip over onto four paws on each developmental day. Negative geotaxis was tested by placing each pup, facing downwards, on a screen angled at 45° from parallel, and measuring the time for it to completely turn and to climb to the top of the screen. Failures to turn and climb were recorded as a maximum score of 30-s. Circle transverse was tested by placing each pup in the center of a circle with a 5″ (12.5 cm) diameter drawn on a laminated sheet of 8.5″ x 11″ white paper, and measuring the time for it to exit the circle. Failures to exit the circle were recorded as a maximum score of 30-s.

### Elevated-plus maze

The assay was performed using a mouse EPM (model ENV-560A) purchased from Med Associates (St. Albans, VT) and performed as previously described^70^.

### Light↔dark conflict

The light↔dark assay was performed in accordance with previously described procedures^70^. The mouse was allowed to explore freely for 10-min. Time in the dark side chamber and total number of transitions between the light and dark side chambers were automatically recorded during the 10-min session.

### Open Field

General exploratory locomotion in a novel open field arena was evaluated as previously described^46,64,70^. Total distance traversed, horizontal activity, vertical activity, and time spent in the center were automatically measured to assess gross motor abilities in mice. Repeated-measures ANOVA was used to detect differences in horizontal, vertical, total, and center time activity obtained during the open field assay. Multiple comparisons were corrected for using Sidak post hoc methods and F, degrees of freedom, and p-values are reported.

### Spontaneous Alternation in a Y-maze

Spontaneous alternation was assayed using methods modified based from previous studies^64^ in mice. One-way ANOVA was used to detect differences in alternation. Multiple comparisons were corrected for using Sidak post hoc methods and F, degrees of freedom, and p-values are reported.

### Novel Object Recognition

The novel object recognition test was conducted as previously described^46,47,71^. The assay consisted of four sessions: a 30-min habituation session, a second 10-min habituation phase, a 10-min familiarization session, and a 5-min recognition test. Sniffing was defined as head facing the object with the nose point within 2 cm from the object. Time spent sniffing each object was scored by an investigator blind to both genotype and treatment. Recognition memory was evaluated by time spent sniffing the novel object versus the familiar object and innate side bias was accounted for by comparing sniff time of the two identical objects during familiarization. Within genotype repeated-measures ANOVA was used to analyze novel object recognition using novel versus familiar objects as comparison. F, degrees of freedom, and p-values are reported.

### Balance beam walking

Balance beam walking is a standard measure of motor coordination and balance^72,73^. We followed a procedure similar to methods previously described using our behavioral core^74^. Latency to traverse the length of the beam, number of footslips off the edge of the beam, and falls (if any), are scored by the investigator from coded video recordings. Approximately four trials per day for three days represents a standard training protocol.

### Rotarod

Motor coordination, balance, and motor learning were assessed using an accelerating rotarod (Ugo Basile, Schwenksville, PA) as previously described^75,76^. The task requires the mice to walk forward in order to remain on top of the rotating cylinder rod. Mice were given 3 trials per day with a 30–60-minute intertrial rest interval. Mice were tested over two consecutive days for a total of 6 trials. Latency to fall was recorded with a 300-second maximum latency.

### Repetitive self-grooming

Spontaneous repetitive self-grooming behavior was scored as previously described^64,70,75, 77, 78^. Each mouse was placed individually into a standard mouse cage (46 cm long × 23.5 cm wide × 20 cm high). The first 10-min was habituation and was unscored. Each subject was scored for cumulative time spent grooming all the body regions during the second 10 min of the test session.

### Three chambered social approach

Social approach was tested in an automated three-chambered apparatus using methods similar to those previously described^46,64,70,78,79^. Three zones, defined using the EthoVision XT software, detected time in each chamber for each phase of the assay. Direction of the head, facing toward the cup enclosure, defined sniff time. The subject mouse was first contained in the center chamber for 10 min, then allowed to explore all three empty chambers during a 10 min habituation session, then allowed to explore the three chambers containing a novel object in one side chamber and a novel mouse in the other side chamber. Lack of innate side preference was confirmed during the initial 10 min of habituation to the entire arena. Novel stimulus mice were 129Sv/ImJ, a relatively inactive strain, aged 10–14 weeks, and matched to the subject mice by sex. Number of entries into the side chambers served as a within-task control for levels of general exploratory locomotion.

### Male-female social interaction

The male–female reciprocal social interaction test was conducted as previously described^64,70,75,77,78^. Briefly, each freely moving male subject was paired for 5-min with a freely moving unfamiliar estrous WT female. Duration of nose-to-nose sniffing, nose-to-anogenital sniffing and following were scored using Noldus Observer 8.0XT event recording software (Noldus, Leesburg, VA). Ultrasonic vocalization spectrograms were displayed using Avisoft software and calls were identified manually by a highly trained investigator blinded to genotype.

### Fear conditioning

Delay contextual and cued fear conditioning was conducted using an automated fear-conditioning chamber (Med Associates, St Albans, VT, USA) as previously described^46,64,70^. Training consisted of a 2-min acclimation period followed by three tone–shock (CS–US) pairings (80-dB tone, duration 30 s; 0.5-mA footshock, duration 1 s; intershock interval, 90 s) and a 2.5-min period during which no stimuli were presented. The environment was well lit (∼100 lx), with a stainless steel grid floor and swabbed with vanilla odor cue (prepared from vanilla extract; McCormick; 1:100 dilution). A 5-min test of contextual fear conditioning was performed 24 h after training, in the absence of the tone and footshock but in the presence of 100 lx overhead lighting, vanilla odor and chamber cues identical to those used on the training day. Cued fear conditioning, conducted 48 h after training, was assessed in a novel environment with distinct visual, tactile and olfactory cues. The cued test consisted of a 3-min acclimation period followed by a 3-min presentation of the tone CS and a 90-s exploration period. Cumulative time spent freezing in each condition was quantified by VideoFreeze software (Med Associates).

### Acoustic Startle and Prepulse inhibition

Subjects were tested in San Diego Instruments startle chambers using standard methods as described previously^75, 80^. One trial type measured the response to no stimulus (baseline movement). The other five trial types measured startle responses to 40 ms sound bursts of 80, 90, 100, 110, or 120 dB. The maximum startle amplitude over this sampling period was taken as the dependent variable. For prepulse inhibition of acoustic startle, mice were presented with each of seven trial types across six discrete blocks of trials for a total of 42 trials, over 10.5 min. One trial type measured the response to no stimulus (baseline movement) and another measured the startle response to a 40 ms, 110 dB sound burst. The other five trial types were acoustic prepulse stimulus plus acoustic startle stimulus trials. The trial types were presented in pseudorandom order such that each trial type was presented once within a block. Prepulse stimuli were 20 ms tones of 74, 78, 82, 86, and 92 dB intensity, presented 100 ms before the 110 dB startle stimulus. The maximum startle amplitude over this sampling period was taken as the dependent variable. A background noise level of 70 dB was maintained over the duration of the test session.

### Seizure Susceptibility Following Administration of Pentylenetetrazole

Subjects were weighed then administered pentylenetetrazole (80 mg/kg) intraperitoneally. Dosing was conducted in the morning (9:00-12:00) in a dim (∼30 lux) empty holding room. Directly after administration of the convulsant, subjects were placed in a clean, empty cage and subsequent seizure stages were live-scored for 30-min. Seizure stages were scored using latencies to (1) first jerk/Straub’s tail, (2) loss of righting, (3) generalized clonic-tonic seizure, and (4) death. Time to each stage was taken in seconds and compared by genotype. Unpaired Student’s t-tests were used to analyze latencies to first jerk, loss of righting, generalized clonic-tonic seizure, and death.

### Thermal Induction of Seizures in Juveniles [Febrile Seizure Paradigm]

Subjects were acclimated to an arena for 10 mins maintaining temperature at 37.5 degrees. Temperature was increased 0.5 degrees every 2 mins until animal showed behavioral seizure or a max temperature of 42.5 degrees was reached.

Observer recorded the type of seizure, and the temperature at which the behavioral seizure was observed using the Racine score from the literature for Febrile Seizures:

1. staring
2. head nodding
3. unilateral forelimb clonus
4. bilateral forelimb clonus
5. rearing and falling
6. clonic seizure

### EEG Implantation

Wireless EEG transmitters were implanted in anesthetized test animals using continuous isoflurane (2-4%). A 2-3 cm midline incision was made over the skull and trapezius muscles, then expanded to expose the subcutaneous space. Implants were placed in the subcutaneous pocket lateral to the spine to avoid discomfort of the animal and displacement due to movement. Attached to the implant were 4 biopotential leads made of a Nickel-Colbalt based alloy insulated in medical-grade silicone, making up two channels that included a signal and reference lead. These leads were threaded towards the cranial part of the incisions for EEG and EMG placement. The periosteum was cleaned from the skull using a sterile cotton-tip applicator and scalpel then two 1mm diameter burr holes were drilled (1.0mm anterior and 1.0mm lateral; −3.0mm posterior and 1.0mm lateral) relative to bregma. This lead placement allowed for measurement of EEG activity across the frontal cortical area. Steel surgical screws were placed in the burr holes and the biopotential leads were attached by removing the end of the silicone covering and tying the lead to its respective screw. Once in place, the skull screws and lead connections were secured using dental cement. For EMG lead placement, the trapezius muscles of the animal were exposed, and each lead was looped through and sutured to prevent displacement. Finally, the incision was sutured using non-resorbable suture material and the animals were placed in a heated recovery cage where they received Carpofen (5mg/kg; i.p.) directly after surgery and 24 hours post-surgery as an analgesic. Subjects were individually caged with *ad libitum* access to food and water for 1-week before EEG acquisition and monitored daily to ensure proper incision healing and recovery. Each implantation surgery took <45-min and no fatalities were observed.

### EEG acquisition

EEG data was acquired using Ponemah (Data Sciences International, St. Paul, MN, USA) and subsequently analyzed using the Neuroscore automated software (Data Sciences International, St. Paul, MN, USA). Subjects were recorded for 24-h baseline in their home cage before administration of 80 mg/kg pentylenetetrazole (Sigma Aldrich, St. Louis, MO, USA) injected intraperitoneally. EEG and EMG were continuously sampled at 500 Hz. Spiking was defined as activity above an absolute threshold of 200 µV-1000 µV that lasted between 0.5 and 100 ms, while spike trains had a minimum duration of 0.5s, a minimum spike interval of 0.05s and a minimum of 4 consecutive spikes. Power spectral densities were determined using a periodogram transformation from amplitude to frequency domains then log transformed for clearer data illustration. Latency to seizure onset and subsequent death following administration of PTZ was first quantified by observed latencies then confirmed by spectral EEG and amplitude response read-outs. One-way ANOVA was used to analyze bouts of spike train activity and latency to seizure onset and death between genotypes. An overall ANOVA was used to detect a genotype difference across power bands, then genotype differences were analyzed within power bands using multiple comparisons.

### Luciferase assay

We constructed luciferase reporter plasmids by cloning an ∼900 bp region containing human 1b^30^ into the pGL4.24 vector (Promega) upstream of the minP, primers in Table 4. HEK293 cells or SK-N-SH cells (40%–60% confluent) were transfected using Lipofectamine 3000 with each construct (400 ng) and the Renilla luciferase expression vector pRL-TK (40 ng; Promega) in triplicate. After 24 hours, the luciferase activity in the cell lysates was determined using the Dual Luciferase Reporter System (Promega). Firefly luciferase activities were normalized to that of Renilla luciferase, and expression relative to the activity of an inactive region of noncoding DNA (NEG2) was noted.

**Table 4:**
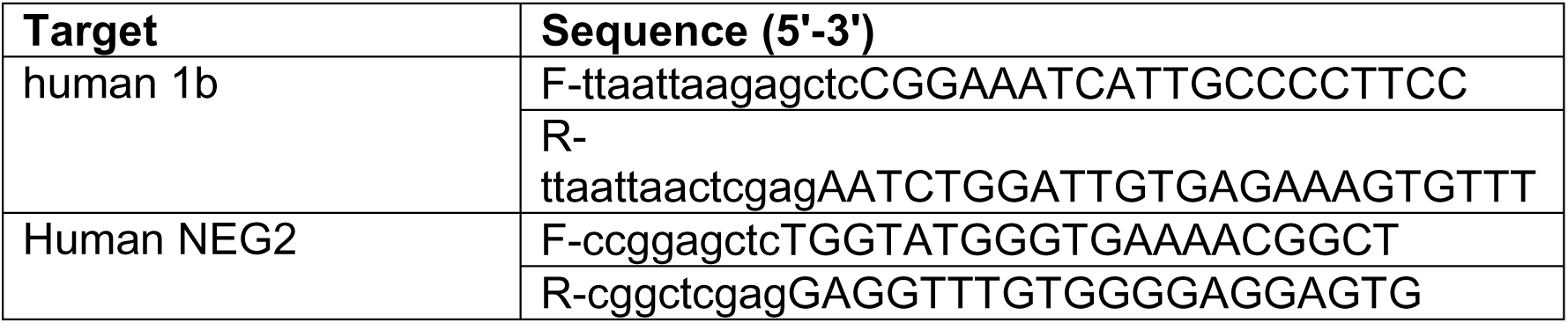
Primers for cloning regions from human DNA.

### CRISPR/dCas9 in HEK293 cells

HEK293 cells were transfected with 500 ng of equimolar pooled SCN1A_h1b sgRNAs (Table 4) and 500 ng dCas9^p300Core^ (Addgene, plasmid #61357) using Lipofectamine 3000. After 24 hours media was refreshed. 48 hours following transfection RNA was collected using RNAqueous kit (Ambion) and cDNA was generated using Superscript III reverse transcriptase (Invitrogen). Changes in gene expression were quantified via qPCR using SYBR green, primers are listed in Table 2.

**Table 4:**
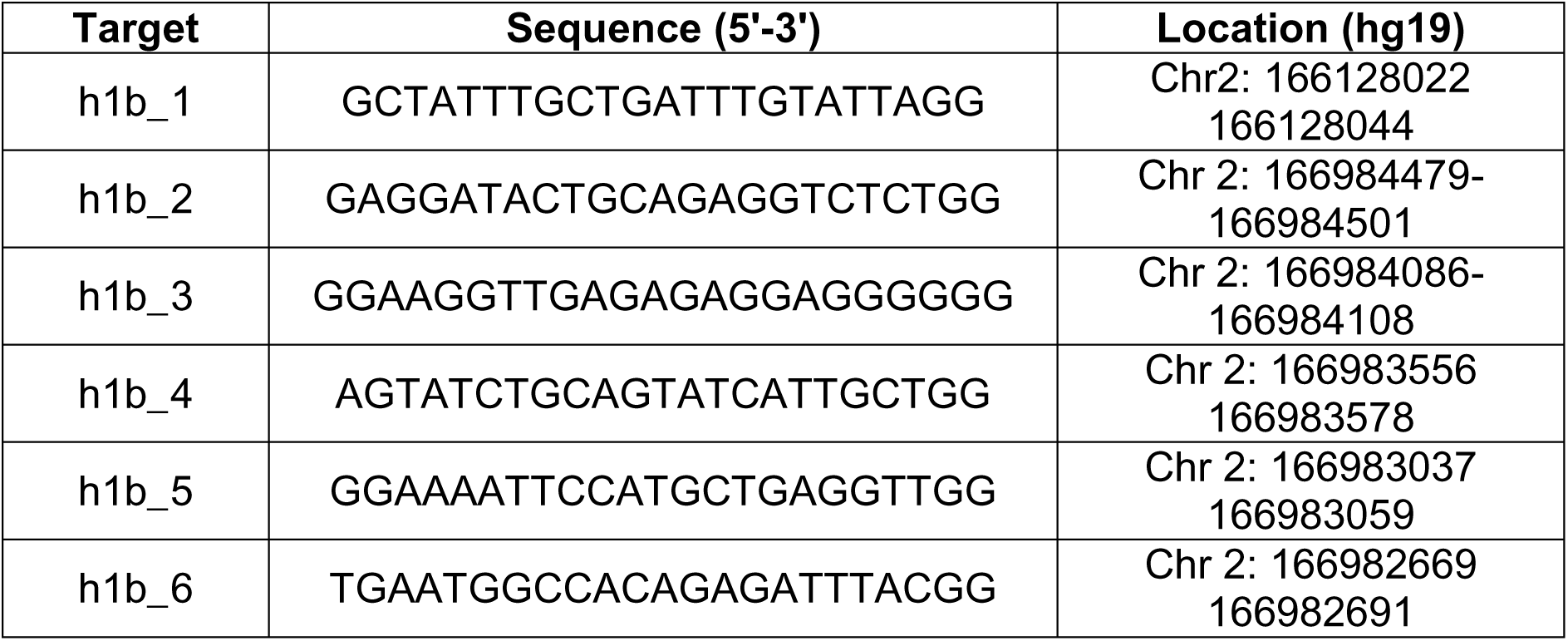
sgRNA sequences for CRISPR dCas9 induction.

